# Emergence of Functional Heart-Brain Circuits in a Vertebrate

**DOI:** 10.1101/2025.09.22.677693

**Authors:** L. Hernandez-Nunez, J. Avrami, S. Shi, A. Markarian, A. Kim, J. Boulanger-Weill, V. Ruetten, A. Zarghani-Shiraz, M. Ahrens, F. Engert, M.C. Fishman

## Abstract

The early formation of sensorimotor circuits is essential for survival. While the development and function of exteroceptive circuits and their associated motor pathways are well characterized, far less is known about the circuits that convey viscerosensory inputs to the brain and transmit visceromotor commands from the central nervous system to internal organs. Technical limitations, such as the *in utero* development of viscerosensory and visceromotor circuits and the invasiveness of procedures required to access them, have hindered studies of their functional development in mammals. Using larval zebrafish—which are genetically accessible and optically transparent—we tracked, *in vivo*, how cardiosensory and cardiomotor neural circuits assemble and begin to function. We uncovered a staged program. First, a minimal efferent circuit suffices for heart-rate control: direct brain-to-heart vagal motor innervation is required, intracardiac neurons are not, and heart rate is governed exclusively by the motor vagus nerve. Within the hindbrain, we functionally localize a cholinergic vagal premotor locus that engages this early efferent control. Second, sympathetic innervation arrives and enhances the dynamics and amplitude of cardiac responses, as neurons in the most anterior sympathetic ganglia acquire the ability to drive cardiac acceleration. These neurons exhibit proportional, integral, and derivative–like relationships to heart rate, consistent with controller motifs that shape gain and dynamics. Third, vagal sensory neurons innervate the heart. Distinct subsets increase activity when heart rate falls or rises, and across spontaneous fluctuations, responses to aversive stimuli, and optogenetically evoked cardiac perturbations, their dynamics are captured by a single canonical temporal kernel with neuron-specific phase offsets, supporting a population code for heart rate. This temporally segregated maturation isolates three experimentally tractable regimes—unidirectional brain-to-heart communication, dual efferent control, and closed-loop control after sensory feedback engages—providing a framework for mechanistic dissection of organism-wide heart–brain circuits.

## Introduction

Animals execute sophisticated behavioral escape responses to avoid environmental threats. These movements are coordinated by specific sensorimotor circuits, which receive inputs via specialized sensory neurons that are tuned to respond to environmental cues of ethological relevance. Visual pathways, for example, are tuned to detect the motion or change in size of nearby objects [1], and downstream motor circuits coordinate action sequences that serve to maintain a balance between responding with sufficient strength to achieve the desired goal and avoiding an overreaction that would exhaust the animal and its metabolic reserves [2]. Critically, these sensorimotor processes must be in place and ready to function the moment the animal engages with the world, which in oviparous species can occur very early in development.

Animal survival also depends on sophisticated internal organ responses to environmental threats. For example, cardiac output varies in response to environmental challenges, allowing adaptive oxygen delivery during vigorous behaviors. This regulation occurs at sub-second timescales and requires the activation of neural circuits, which enable fast and bidirectional communication between the brain and internal organs [3,4]. However, the functional development of these circuits remains poorly understood, largely due to technical limitations. In rodents, for instance, monitoring or manipulating heart-brain circuits often requires invasive procedures performed under anesthesia, which disrupts the very homeostatic responses being studied [5,6]. Moreover, in most mammals, including rodents and humans, autonomic innervation of the heart and other viscera begins *in utero*, precluding longitudinal functional imaging during the initial establishment of heart-brain connectivity [7]. An exception is found in marsupials, where autonomic innervation occurs postnatally [8], offering rare developmental access, but these species are experimentally challenging. Thus, a model that combines *ex utero* development, optical accessibility, and intact physiology is ideal to understand how autonomic circuits emerge and function during early life.

Larval zebrafish offer, for several reasons, an ideal alternative to studying the functional development of heart- brain neural circuits in intact animals. First, zebrafish larvae are small, relatively transparent, and amenable to developmental, behavioral, and physiological studies using only modest restraint. Second, the structure of the neural circuits that connect internal organs and the brain is conserved in bony fish, reptiles, birds, and mammals [9]. In all these vertebrates, organ regulation is carried out by the motor vagus nerve and the sympathetic system, which send critical control signals from the brain and the spinal cord to the organs.

Similarly, in those vertebrates, most viscerosensory signals are carried by the sensory vagus nerve to the brain [10]. Third, cardiac responses can be measured and manipulated in larval zebrafish with optical methods.

These advantages are crucial but not sufficient; we therefore developed hardware and software tools, as well as data analysis pipelines and transgenic animals that allow us to study the functional development of heart- brain motor and sensory circuits.

The heart is intrinsically autorhythmic and begins beating before cardiac innervation is established. In mammals, autonomic projections reach the heart during prenatal development, and physiological studies indicate that parasympathetic influence emerges early while sympathetic regulation starts later in gestation [7,36]. Much less is known about the developmental establishment of cardiac sensory pathways or when afferent activity first begins to represent cardiac state. More broadly, existing studies have not resolved how neural activity, cardiac regulation, and anatomical connectivity emerge together during the initial assembly of heart–brain circuits.

In zebrafish, anatomical studies have characterized the intrinsic and extrinsic innervation of the mature heart, and stimulation of the adult motor vagus and sympathetic nerves produces bradycardia and tachycardia, respectively [11,12]. Moreover, *in vitro* preparations of the zebrafish heart have been developed to study the effects of pharmacological agents, including vapor anesthetics, on neural regulation of cardiac function [13]. Experiments with adult zebrafish, however, have some of the same technical limitations as experiments with rodents, including a lack of optical access, the need for invasive procedures, and the inability to concurrently observe autonomic neural dynamics. Here, we establish larval zebrafish as a model for heart-brain interactions by characterizing the function of motor and sensory circuits of the heart during development and identifying the earliest stages at which sensorimotor control of the heart can be studied while retaining optical access to both the heart and brain.

Using custom instrumentation that combines calcium imaging, optogenetics, and quantitative all-optical cardiac physiology, we resolve how heart–brain circuits acquire functional control and structured relationships between neural activity and cardiac dynamics during development. Early in development, a minimal efferent circuit is sufficient for neural modulation of heart rate: direct brain-to-heart vagal projections provide the first detected efferent cardiac pathway, before intracardiac neurons and sympathetic innervation are established. Within the hindbrain, we functionally localize a cholinergic vagal premotor locus that engages this early efferent control.

As development proceeds, sympathetic innervation arrives and enhances the amplitude and kinetics of cardiac responses. Neurons in the most anterior sympathetic ganglia acquire the capacity to accelerate the heart and exhibit proportional, integral, and derivative-like relationships to heart rate, indicating controller motifs that shape gain and dynamics. Soon after, vagal sensory neurons (VSNs) innervate the heart. Distinct VSN subsets increase activity when heart rate falls or rises. These VSNs encode heart rate during spontaneous fluctuations, responses to aversive stimuli, and optogenetically evoked cardiac perturbations. In all of those cases, their dynamics are captured by one temporal basis function whose phase-shifted instances tile heart rate trajectories, consistent with a population code for heart rate.

Our findings uncover how the vertebrate nervous system progressively assembles a closed-loop control architecture for cardiac regulation, beginning with descending motor control and culminating in the engagement of sensory feedback. Human autonomic disorders, such as postural orthostatic tachycardia syndrome (POTS) and long COVID-associated dysautonomia, are hypothesized to involve disruptions in feedback control of the heart, including altered sensory gain and defective reflex adaptation [14, 45]. The framework we present enables *in vivo* testing of these hypotheses at cellular resolution by allowing precise manipulation of individual system components and direct observation of resulting deficits in cardiac dynamics.

## Results

### The Emergence of a Minimal Functional Circuit for Neurocardiac Control

The two-chambered heart of the zebrafish embryo starts beating 24 hours post-fertilization, with atrial contraction followed by ventricular contraction, driven by its intrinsic pacemaker [16, 17]. To investigate the emergence of a minimal circuit for neurocardiac control in larval zebrafish, we developed a system, as shown in Figure 1A, that enables the quantification of the heart’s physiological function while simultaneously exposing the animal to visual environmental stimuli. In our setup, the larval zebrafish head is immobilized using agarose, while the tail is free to move (Supp. Fig. 1G). We quantified individual atrial or ventricular contractions based on the transmission of near-infrared light, as shown in Figure 1B. Using this approach, we examined the spontaneous heart rate of fish during development and found, consistent with prior work [18, 19], that basal heart rate first decreases from 4 to 5 days post fertilization (dpf) and increases from 5 to 7 dpf (Supp. Fig. 1B).

**Figure 1.**
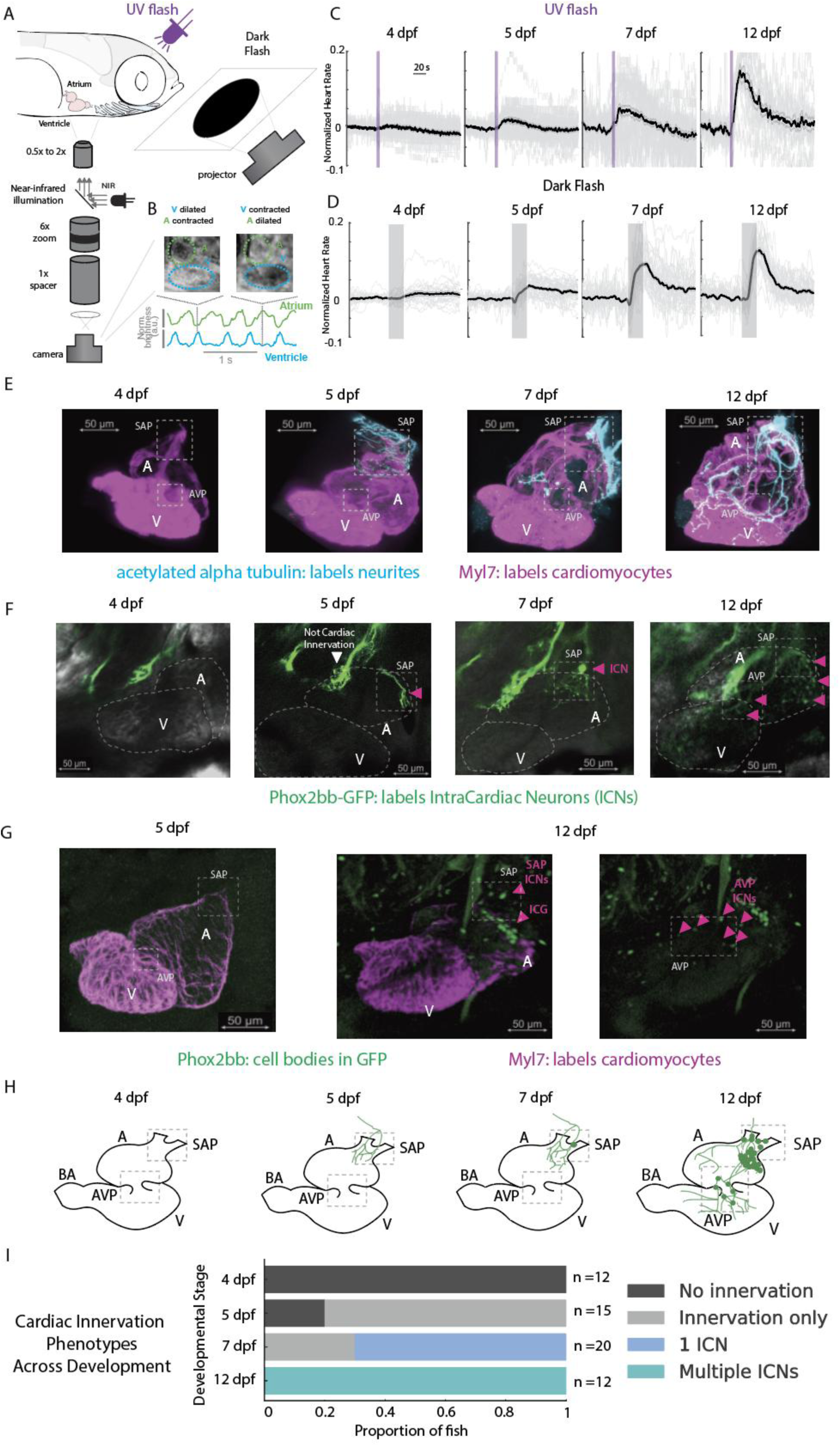
Developmental tracking of the intracardiac nervous system reveals a minimal structure for neural control of the heart. **(A)** Schematic representation of a custom-built experimental setup for imaging the visceral cavity of zebrafish larvae, including the heart. Visual stimuli can be delivered with a projector or an LED. **(B)** HR extraction. Top: example NIR frames during ventricular dilation and contraction. In transmitted NIR images, the atrium and ventricle appear brighter during dilation, when optical density along the imaging path is lower, and darker during contraction, when the chamber contents are compressed and transmit less light. Bottom: NIR intensity traces from the atrium (A, green) and ventricle (V, cyan) used to compute beat timing (1 s scale bar). **(C)** HR responses to UV flash across development (4, 5, 7, 12 dpf). Thick black line, mean normalized HR; gray shading, ± SE; vertical violet bars, stimulus onset; n=14–20 fish per age. **(D)** HR responses to dark flash across the same ages. Plotting as in (C) with gray stimulus bars. **(E)** Cardiac innervation timeline in fixed hearts stained for acetylated α-tubulin (neurites, cyan) and Myl7 (myocardium, magenta). Innervation appears at SAP by 5 dpf, extends through the atrium by 7 dpf, and reaches AVP and ventricle by 12 dpf. Scale bars, 50 µm. **(F)** *In vivo* phox2bb:GFP imaging of the heart region. 4–5 dpf: axons approach the heart but no ICNs are present; 7 dpf: first ICN appears; 12 dpf: multiple ICNs (magenta arrowheads) are evident. Scale bars, 50 µm. **(G)** Immunohistochemistry for Phox2bb (cell bodies, green) and Myl7 (magenta). 5 dpf: no ICN somata in the heart; 12 dpf: ICNs near SAP (≈6), near AVP (≈6), and a dorsal-posterior atrial cluster (ICG; ≈16). Scale bars, 50 µm. **(H)** Schematic summary of the developmental sequence: 5 dpf—axon entry at SAP; 7 dpf—first ICN; 12 dpf— ICNs at SAP/AVP and ICG. BA, bulbus arteriosus. **(I)** Population summary of innervation phenotypes by age (proportion of fish; 4 dpf n=12, 5 dpf n=15, 7 dpf n=20, 12 dpf n=12). Categories: no innervation; innervation only; ≥1 ICN; multiple ICNs. **Conventions.** HR traces are normalized per fish. SE shading indicates standard error (between-fish variability). All scale bars, 50 µm. Abbreviations: NIR, near-infrared; SAP, sinoatrial plexus; AVP, atrioventricular plexus; ICN, intracardiac neuron; ICG, intracardiac ganglion; BA, bulbus arteriosus.

To induce transient changes in heart rate, we used a brief UV flash (Fig. 1C), a flash of darkness (Fig. 1D), or looming stimuli (Supp. Fig. 1C-F), which are well-studied visual threats. We subjected larval zebrafish to these stimuli across developmental stages ranging from 4 to 12 dpf. We found that while at 4 dpf, fish are capable of behavioral escapes as marked by tail flicks (Supp. Fig. 1G), their heart rate showed no detectable stimulus-evoked modulation (Fig. 1C, D, Supp. Fig. 1C, I). At 5 dpf, larval fish start displaying increased heart rate responses to UV flash, dark flash, and looming (Fig. 1C, D, Supp. Fig. 1D, I). These responses change their temporal dynamics and intensity as the fish matures. By 12 dpf, the response is larger, more rapid in onset and adaptation, and more stereotypical than at earlier stages (Fig. 1C, D, Supp. Fig. 1F, I).

Baseline interbeat intervals became more variable across development. We quantified short-timescale variability during pre-stimulus periods using the root mean square of successive differences (RMSSD) between consecutive interbeat intervals, normalized to the mean interbeat interval. Normalized RMSSD increased 2.66- fold between 4 and 5 dpf (bootstrap 95% CI, 2.04–3.27) and remained elevated at 7 and 12 dpf (Supp. Fig. 2A, B). The coefficient of variation of interbeat intervals (CVNN) showed a similar developmental pattern.

Moreover, the relationship between absolute RMSSD and mean interbeat interval differed across ages, indicating that the developmental increase in variability could not be explained solely by differences in mean heart rate (Supp. Fig. 2C, D).

To define the minimal neural structure required for heart rate modulation, we examined the spatiotemporal development of cardiac innervation using immunostaining for acetylated alpha-tubulin to label neurites and Myl7 to label the myocardium (Fig. 1E). We observed that cardiac innervation begins at 5 days post-fertilization (dpf), when the first neurites reach the entry of the atrium, known as the sinoatrial plexus (SAP), a region that includes the pacemaker cells [20,21]. As development proceeds, innervation expands in a stereotyped sequence: the atrium is innervated by 7 dpf, followed by the atrioventricular plexus (AVP) near the valve region, and the ventricle by 12 dpf.

Next, we asked whether intracardiac neurons are required for the initiation of heart rate control. To track their development, we used transgenic zebrafish expressing GFP under the control of the phox2bb promoter, which labels all intracardiac neurons (ICNs) [47]. Consistent with our acetylated α-tubulin staining, we observed no cardiac innervation at 4 dpf (Fig. 1F). By 5 dpf, neurites begin to reach the heart, but no intracardiac neurons are yet present (Fig. 1F; Supp. Vid. 1). The first intracardiac neuron migrates into the heart by 7 dpf (Fig. 1F; Supp. Vid. 2), and by 12 dpf, multiple intracardiac neurons are evident (Fig. 1F; Supp. Vid. 3). To visualize neuronal somata more clearly, we performed immunostaining for Phox2bb alongside Myl7 to label the myocardium (Fig. 1G). At 5 dpf, no neuronal cell bodies were observed within the heart. In a representative 12 dpf preparation, we identified approximately 6 neurons near the SAP, 6 near the AVP, and a dense dorsal-posterior atrial cluster of approximately 16 neurons, which we designate the Intracardiac Ganglion (ICG).

Because somata within the ICG frequently overlap, these values describe the representative preparation rather than an exhaustive population count across animals.

To summarize the timing and variability of cardiac innervation and ICN emergence, we quantified developmental progression across animals (Fig. 1H, I): at 4 dpf, 0/12 fish showed cardiac innervation; at 5 dpf, 12/15 fish showed innervation; at 7 dpf, 14/20 fish had at least one intracardiac neuron; and by 12 dpf, all 12/12 fish displayed robust innervation of both cardiac chambers with multiple intracardiac neurons (Supp. Fig. 1H). Together, these results show that functional cardiac innervation precedes the arrival of intracardiac neurons, establishing that early modulation of heart rate occurs through direct descending motor pathways alone.

### Formation of the Direct Brain-to-Heart Motor Circuit

We next sought to determine which branches of the autonomic nervous system contribute to progressive innervation of the heart between 5 and 12 dpf. Because the motor vagus nerve is cholinergic [10], we began by labeling it with antibodies against choline acetyltransferase (ChAT), the enzyme responsible for synthesizing acetylcholine, and labeling the myocardium with antibodies against Myl7. This approach allowed us to visualize cholinergic innervation of the heart beginning at 5 dpf (Fig. 2A).

**Figure 2.**
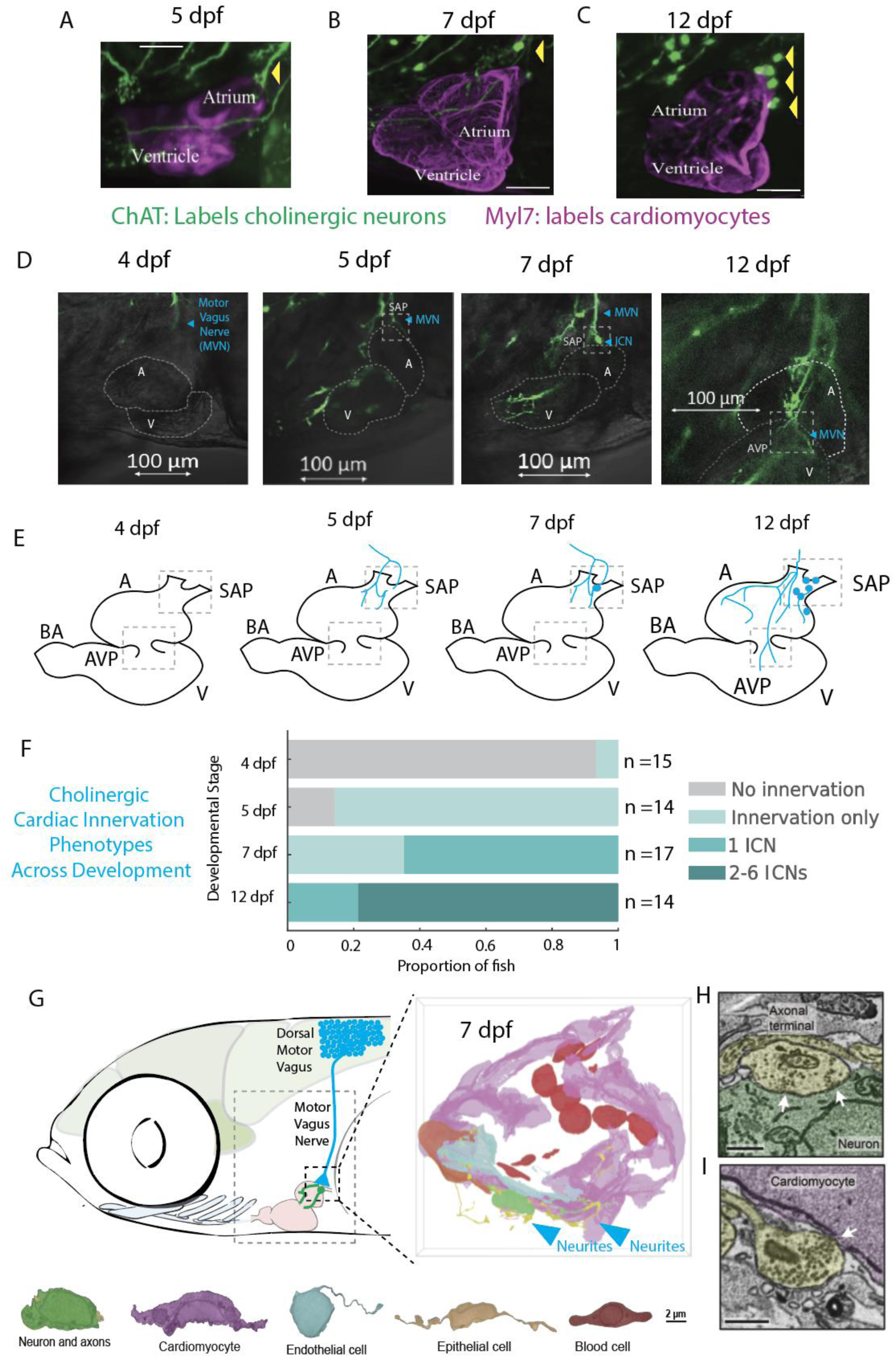
Formation of a direct cholinergic brain-to-heart motor circuit. **(A–C)** Immunostaining for ChAT (green; cholinergic fibers/neurons) and Myl7 (magenta; myocardium) at 5, 7, 12 dpf. Cholinergic fibers reach the sinoatrial region by 5 dpf; the first ChAT-positive ICN appears at 7 dpf; multiple ICNs cluster near the SAP by 12 dpf. Scale bars, 50 µm. **(D)** *In vivo* volumetric imaging in ChaTA-Gal4>UAS-GFP larvae with transmitted-light myocardium. Left→right: 4 dpf (no innervation), 5 dpf (emerging fibers), 7 dpf (first ICN), 12 dpf (extension toward AVP). SAP/AVP and the atrium (A) and ventricle (V) are outlined; arrowheads mark neurites/ICN. Scale bars, 100 µm. **(E)** Schematic progression of cholinergic cardiac innervation across ages: fibers first contact the SAP (5 dpf), a single ICN appears (7 dpf), then 2–6 ICNs are present (12 dpf). BA, bulbus arteriosus. **(F)** Population summary of cholinergic phenotypes by age: no innervation (gray), innervation only (light teal), one ICN (teal), 2–6 ICNs (dark teal). 4 dpf n=15; 5 dpf n=14; 7 dpf n=17; 12 dpf n=14. Among the 12 dpf animals, the number of distinguishable labeled ICNs was 1 neuron in 3 fish, 2 neurons in 5 fish, 3 neurons in 3 fish, and 6 neurons in 3 fish. These values refer to distinguishable labeled somata. **(G)** ssEM reconstruction of the SAP at 7 dpf with segmented cell types (legend). Neurites approaching the SAP are indicated (arrows). Scale bar, 2 µm. **(H)** EM micrograph showing a vesicle-rich axonal terminal contacting the ICN (arrowheads). **(I)** EM micrograph showing a synaptic contact from the ICN onto a cardiomyocyte (arrowheads). Notes. Cholinergic identity of the first ICN was corroborated by HCR-FISH for ChAT and overlap with ChaTA- Gal4>GFP and phox2bb:GFP (Supp. Fig. 3). No ChAT-positive ventricular innervation was detected through 12 dpf. **Acronyms:** A, atrium; V, ventricle; SAP, sinoatrial plexus; AVP, atrioventricular plexus; ICN, intracardiac neuron; BA, bulbus arteriosus.

We also detected ChAT-positive intracardiac neurons (ICNs), including the first neuron to arrive at 7 dpf, suggesting that this cell is a postganglionic neuron of the motor vagus nerve (Fig. 2A–C). Motor vagus postganglionic neurons are known to be cholinergic and receive input from vagal preganglionic fibers and form direct connections with cardiac tissue [23]. By 12 dpf, we observed six ChAT-positive ICNs near the sinoatrial plexus (SAP). We confirmed the cholinergic identity of the first ICN using fluorescent in situ hybridization chain reaction (HCR) for ChAT, together with GFP boosters to confirm overlap with ChaTA-Gal4, UAS-GFP and phox2bb:GFP (Supp. Fig. 3). We did not detect neurons by the atrioventricular plexus (AVP) or elsewhere, suggesting that the other ICNs identified in Fig. 1G may use alternative neurotransmitters.

**Figure 3.**
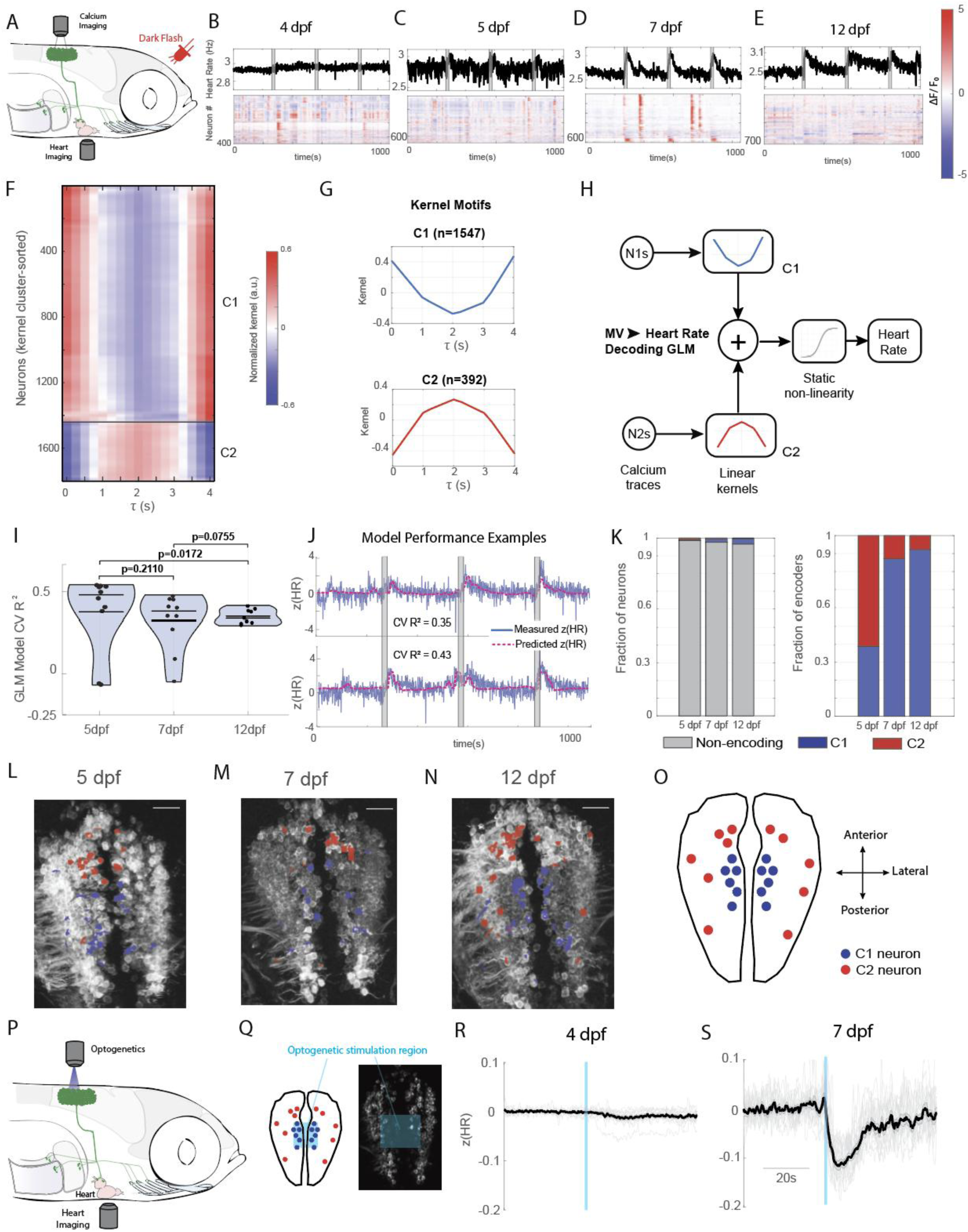
Cholinergic hindbrain decoding of heart rate and emergence of a vagal premotor locus. **(A)** Experimental schematic. Calcium imaging of hindbrain cholinergic neurons (*ChaTA-Gal4*), simultaneous optical heart-rate (HR) readout, and dark-flash stimuli. **(B–E)** Example recordings across development (4, 5, 7, 12 dpf). Top: HR (Hz). Bottom: representative neuronal ΔF/F_0_ heatmaps (rows, neurons; columns, time). **(F)** Decoding kernels for all predictive neurons (exceeding Otsu threshold on cross-validated performance) plotted as a heatmap and ordered by shape similarity. Kernels are causal finite-impulse responses spanning 0–4 s in 0.25 s steps. **(G)** Canonical kernel motifs obtained by clustering all predictive kernels: C1 (negative, blue) and C2 (positive, red). Curves show means with shaded standard error (SE); n values indicate the number of neurons assigned to each motif. **(H)** Decoding model. Each neuron contributes a causal linear kernel; outputs are summed and passed through a static sigmoid nonlinearity to capture HR saturation. **(I)** Model performance by age using blocked 5-fold cross-validation (80% train, 20% test per fold). Points, per- fish CV R²; violins, distributions; horizontal bars, medians. Annotated p values are Wilcoxon rank-sum with BH- FDR correction (Methods). Across 31 fish (5, 7, 12 dpf), only 2 at 5 dpf and 2 at 7 dpf fell below CV R² = 0.20; all others achieved CV R² between 0.25 and 0.60. **(J)** Example predictions for two fish near the cohort median (CV R² = 0.35 and 0.43). Predicted HR closely tracks measured HR and stimulus-locked responses. **(K)** Left: fraction of non-encoding and encoding neurons per age (encoders <5% at each age; prevalence increases at 12 dpf relative to 5 dpf). Right: within encoders, fraction assigned to C1 (negative) or C2 (positive). The negative motif expands from ∼40% at 5 dpf to ∼90% at 12 dpf, consistent with the cardioinhibitory role of motor vagus. **(L–N)** Anatomy maps showing the spatial distribution of encoders at 5, 7, and 12 dpf. Blue, C1 (negative); red, C2 (positive). Encoders cluster medially across ages, with positive encoders more anterior and dispersed, and negative encoders concentrated near the midline. **(O)** Summary schematic of encoder locations across fish: reproducible medial cluster of negative encoders and broader, less consistent distribution of positive encoders. **(P)** Optogenetic testing strategy. ChaTA-Gal4 drives CoChR in cholinergic hindbrain neurons; patterned illumination targets the negative-kernel locus; HR recorded simultaneously. **(Q)** Targeted stimulation region for the premotor locus (example plane). **(R)** At 4 dpf, before anatomical heart innervation is detectable, photoactivation does not measurably change HR. **(S)** At 7 dpf, after innervation appears, the same stimulation produces a robust bradycardia. Traces show mean ± SE ; vertical bars mark stimulus onset.

We then used *ChaTA-Gal4, UAS-GFP* transgenic fish to label motor vagal fibers selectively. Because red fluorophores in the heart interfered with the detection of thin axons, we instead imaged the myocardium in transmitted light using a photodiode-based detector, while capturing green fluorescence in a separate channel. This strategy enabled reliable identification of even fine cholinergic neurites, based on their anatomical position and rhythmic motion during the heartbeat. Consistent with our immunostaining and *phox2bb:GFP* imaging, we observed no innervation at 4dpf (Fig. 2D) (Supp. Vid. 4), emerging innervation at 5 dpf (Fig. 2D, Supp. Vid. 5), emergence of the first ICN at 7 dpf (Fig. 2D, Supp. Vid. 6), and further extension to the AVP by 12 dpf (Fig. 2D, Supp. Vid. 7). We did not detect cholinergic innervation of the ventricle at any stage up to 12 dpf.

To quantify the developmental progression of cholinergic cardiac innervation across individual animals, we scored each fish for the presence of cardiac innervation and the number of cholinergic intracardiac neurons (ICNs) observed at 4, 5, 7, and 12 dpf (Fig. 2E, F). At 4 dpf, the vast majority of fish (14/15) lacked any evidence of cholinergic cardiac innervation. By 5 dpf, most fish (12/14) displayed innervation of the sinoatrial region, but none had ICNs. At 7 dpf, 11 of 17 fish showed a single cholinergic ICN, while the remaining 6 had innervation but no ICNs. By 12 dpf, all fish exhibited robust innervation of the atrium and SAP, with 11 of 14 fish containing between two and six ICNs (Fig. 2F). These data highlight a gradual and stage-specific acquisition of cholinergic innervation and ICNs, with clear stereotypy in the timing of each step in the maturation of the vagal motor circuit.

To investigate if the first ICN already receives neural inputs, we performed serial-section electron microscopy (ssEM) of the SAP in a 7 dpf fish (Fig. 2G). We identified five cell types—cardiomyocytes, epithelial cells, blood cells, endothelial cells, and one neuron, consistent with our light microscopy observations (Fig. 2G). Vesicle- rich synaptic contacts were found both onto the ICN (Fig. 2H) and from the ICN onto cardiomyocytes (Fig. 2H, I), supporting its integration as a functional node in the developing cardiac motor circuit.

### A Cholinergic Vagal Premotor Locus in the Hindbrain Engages Cardiac Inhibition

To explore how this first anatomical evidence of cardiac innervation correlates with the onset of physiological control, we used calcium imaging to measure the activity of cholinergic hindbrain neurons in the region of vagal motor and premotor circuitry. Previous anatomical studies have shown that motor vagus neurons in larval zebrafish are cholinergic [10] and located in the hindbrain [24]; thus, we focused on this population. In order to characterize the dynamics of those neurons, we customized a 2-photon imaging rig with an optical path for simultaneous quantification of cardiac function using near-infrared light and brain functional imaging at single-cell resolution using the 2-photon laser (Fig. 3A). Using ChaTA-Gal4 to drive UAS-GCaMP6s expression, we selectively imaged cholinergic hindbrain neurons while concurrently monitoring heart rate. Dark flash stimulation elicited robust tachycardic responses in 5, 7, and 12 dpf fish (Fig. 3C-E) but not in 4 dpf fish (Fig. 3B), consistent with our results in Fig. 1. Neural responses were diverse and even when large fractions of neurons displayed synchronous increases in activity, those were not consistently accompanied by changes in heart rate (Fig. 3B-E).

We asked whether activity in this cholinergic hindbrain population carries a time-resolved signature of cardiac control. To do so, we fit a generalized linear model (GLM) that decodes heart-rate (HR) dynamics from the population activity, assigning each neuron a causal finite-impulse-response kernel that quantifies the temporal weighting of its activity across preceding time lags. Because HR was calculated from individual cardiac intervals, the modeled target retained the beat-resolved variation present in the cardiac recordings, although the temporal precision recoverable from neural activity was limited by the kinetics and sampling rate of the calcium signals. A static sigmoid nonlinearity captured saturation in HR output. To visualize the temporal motifs, we pooled predictive neurons—those exceeding an Otsu threshold on cross-validated predictive power (Methods, Decoding GLMs)—and plotted their kernels as a shape-sorted heatmap (Fig. 3F). This revealed a single canonical waveform that recurred across cells with opposite polarities: some neurons expressed the positive version and others the negative of the same kernel (Fig. 3F-H).

Model performance was evaluated with blocked 5-fold cross-validation (80% train, 20% test per fold; Methods). Across 31 fish (5, 7, and 12 dpf), only 2 fish at 5 dpf and 2 at 7 dpf fell below CV R² = 0.20; in all others, the model predicted HR trends with CV R² ranging from 0.25 to 0.60 (Fig. 3I). Even for animals near the cohort median (CV R² = 0.35 and 0.43), the predicted traces closely tracked stimulus-evoked HR responses (Fig. 3J). Although predictive neurons (defined by the Otsu threshold on CV performance) comprised <5% of recorded cells at each age, their prevalence tripled at 12 dpf relative to 5 dpf (Fig. 3K). Among encoders, the fraction with negative kernels—the HR-decreasing sign—rose from ∼40% at 5 dpf to ∼90% at 12 dpf, consistent with the cardioinhibitory role of the motor vagus (Fig. 3K). Spatially, encoders were concentrated medially across ages; positive-kernel encoders were more anterior and dispersed, whereas negative-kernel encoders formed a reproducible medial cluster (Fig. 3L–O).

The presence of two opposing kernel classes raised the question of whether decoding depended on their combined activity or whether each class independently retained information about heart-rate dynamics. We therefore retrained the GLM using only C1 or C2 neurons. Both restricted models retained substantial cross-validated predictive performance at 5, 7, and 12 dpf, with no significant differences from the combined C1+C2 model at any age examined (Supp. Fig. 4). Thus, predictive information about heart-rate dynamics is distributed across both kernel classes, with either class alone capturing much of the signal recovered by the combined population.

**Figure 4.**
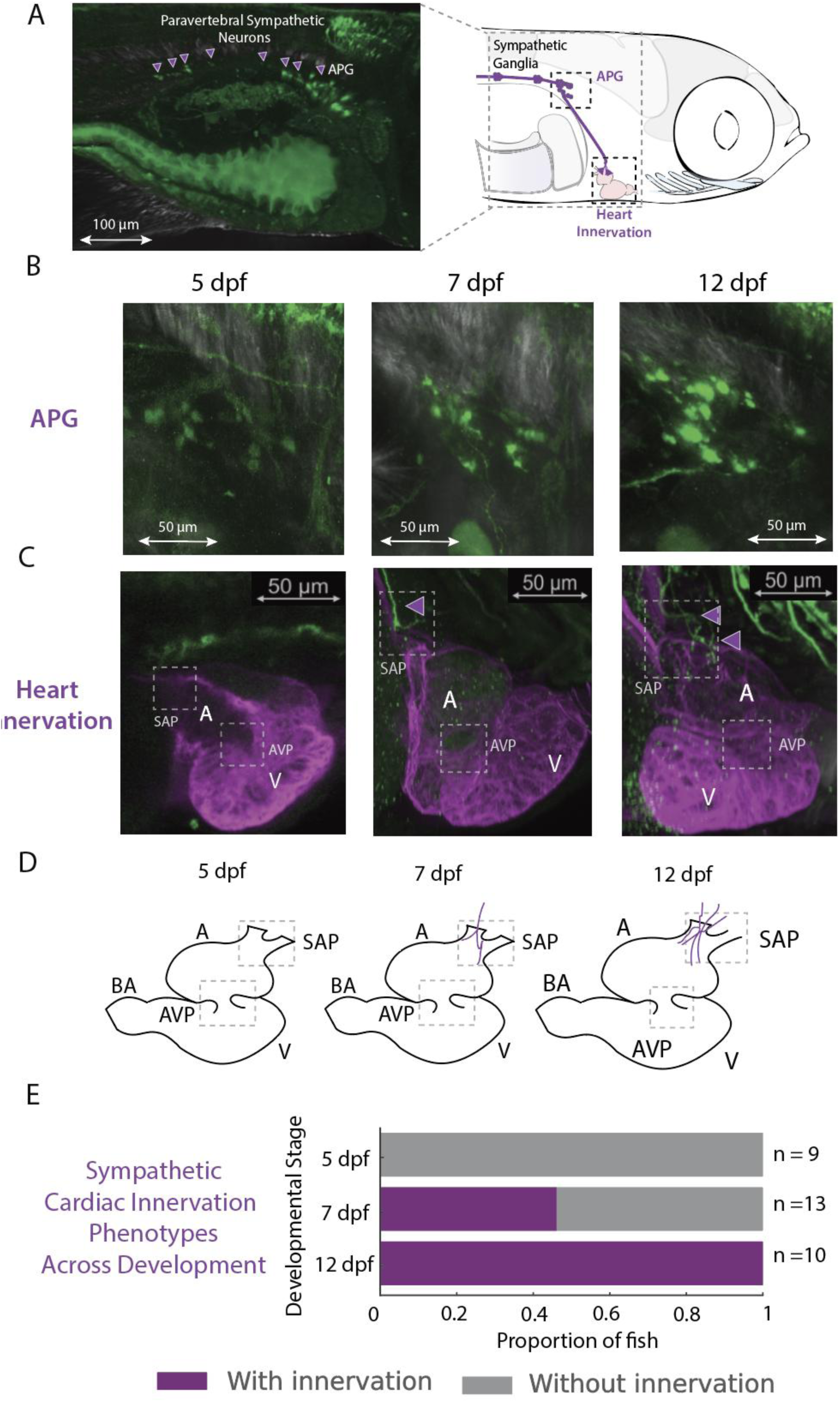
Development of the sympathetic cardiac circuit. **(A)** Paravertebral sympathetic neurons *in vivo*. Arrowheads mark sympathetic neurons along the trunk; APG indicates the anterior paravertebral ganglia that project toward the heart (schematic, right). Scale bar, 100 µm. **(B)** APG morphology over development (5, 7, 12 dpf). TH immunolabeling reveals growth and the characteristic C-shaped configuration. Scale bars, 50 µm. **(C)** Sympathetic cardiac innervation (TH, green) with myocardium (Myl7, magenta). 5 dpf: no cardiac TH fibers. 7 dpf: TH-positive projections reach the SAP (arrowheads). 12 dpf: SAP innervation strengthened; no AVP or ventricular TH fibers detected. Scale bars, 50 µm. **(D)** Schematic summary of innervation patterns at 5, 7, and 12 dpf. BA, bulbus arteriosus; A, atrium; V, ventricle; SAP, sinoatrial plexus; AVP, atrioventricular plexus. **(E)** Population summary of sympathetic cardiac innervation by age (with/without TH-positive fibers at the SAP). 5 dpf n=9: 0% innervated; 7 dpf n=13: 46% innervated; 12 dpf n=10: 100% innervated. **Notes.** TH marks catecholaminergic sympathetic fibers; all panels use TH (green) and Myl7 (magenta) unless indicated. Innervation remained SAP-restricted up to 12 dpf in this cohort.

Given the stereotyped, medial clustering of negative-kernel encoders and the established cardio-inhibitory role of the motor vagus, we hypothesized that activating neurons in this locus would reduce heart rate. We expressed the light-gated channel CoChR under ChaTA-Gal4 in cholinergic hindbrain neurons and delivered patterned photostimulation to the negative-kernel locus using structured illumination steered by dual galvos (Fig. 3P,Q). At 4 dpf, before anatomical heart innervation is detectable, photoactivation of this locus did not measurably alter heart rate (Fig. 3R). By 7 dpf, once innervation is present, the same stimulation decreased heart rate (Fig. 3S). As a control, optogenetic activation of the anterior spinal cord did not result in changes in heart rate (Supp. Fig. 5). These data identify a medial cholinergic hindbrain locus that engages descending vagal cardiac inhibition.

**Figure 5.**
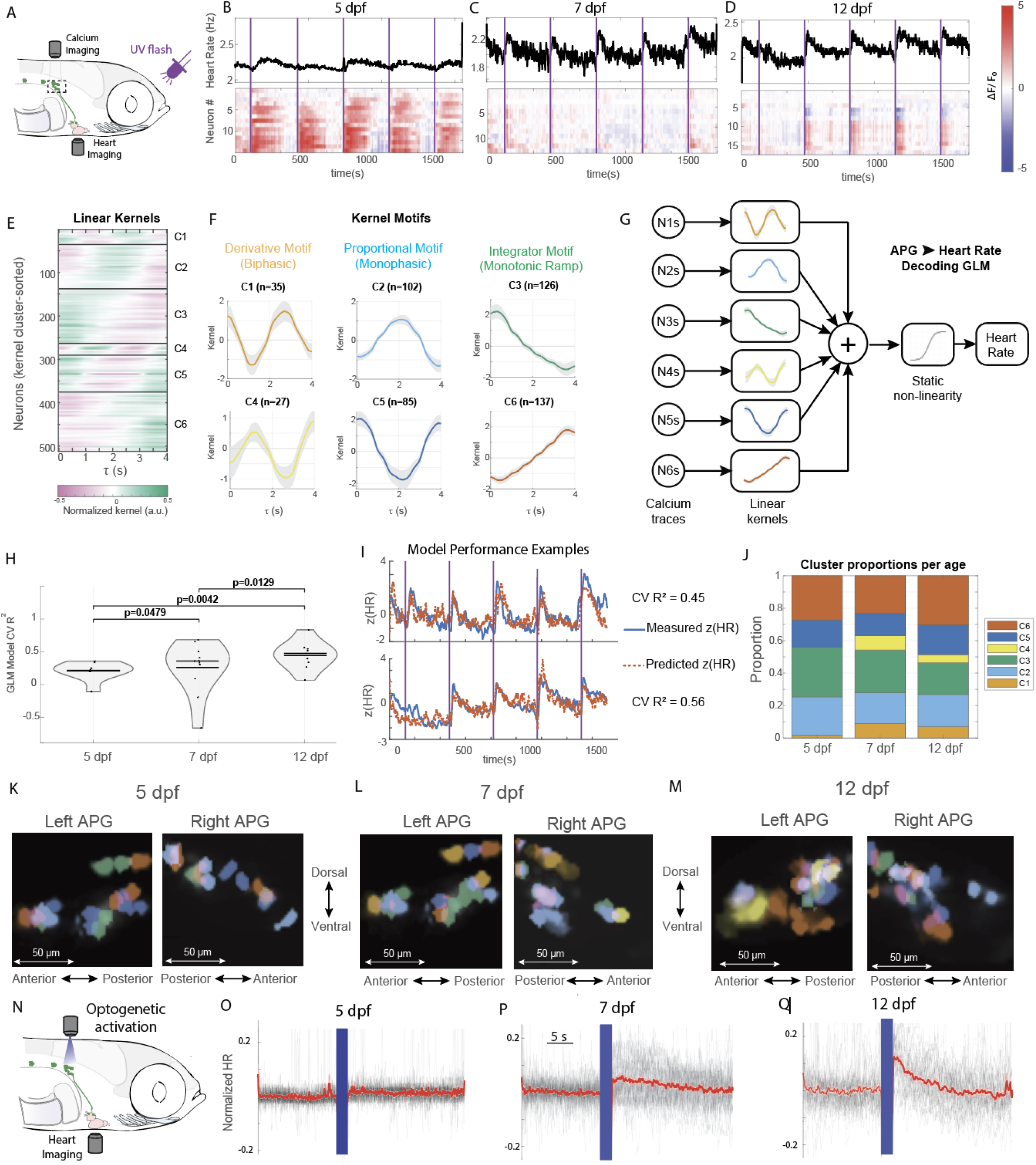
Emergence of control motifs and function in sympathetic cardiac circuits. **(A)** Assay schematic. TH1-Gal4>UAS-GCaMP6f labels sympathetic neurons in the anterior paravertebral ganglia (APG). Calcium activity and heart rate (HR) are recorded simultaneously during brief UV flashes. **(B–D)** Representative recordings at 5, 7, and 12 dpf. Top: HR (Hz). Bottom: APG population activity (ΔF/F heatmaps; rows = neurons; columns = time). Vertical purple bars indicate UV flashes. **(E)** Linear kernels for all predictive neurons, pooled across fish and ordered by shape similarity. Lags: 0–4 s in 0.1-s steps; color denotes normalized kernel amplitude. **(F)** Canonical kernel motifs (cluster means ± SE). Unsupervised clustering reproducibly yielded six motifs: two biphasic/derivative-like (C1, C4), two monophasic/proportional-like (C2, C5), and two monotonic ramp/integrator-like (C3, C6). n per motif indicated. **(G)** Decoding GLM diagram. Each neuron contributes a causal kernel; outputs are summed and passed through a static sigmoid nonlinearity to predict HR. **(H)** Cross-validated performance by age (blocked 5-fold; 80% train/20% test per fold). Points, fish; violins, distributions; bars, medians. P values (above) are Wilcoxon rank-sum with BH-FDR correction. **(I)** Example held-out predictions from the GLM for two fish (CV R² = 0.45 and 0.56). Blue, measured z(HR); orange dashed, model prediction. **(J)** Cluster composition across development. Stacked bars show the proportion of neurons assigned to each motif (C1–C6) per age. Derivative-like motifs are rare at 5 dpf and emerge by 7–12 dpf. **(K–M)** Spatial organization of APG neurons colored by cluster identity in left and right ganglia at 5, 7, 12 dpf. Dorsal–ventral and anterior–posterior axes indicated; scale bars, 50 µm. Neuron positions vary between fish and between sides. **(N)** Optogenetic test of function. TH1-Gal4>UAS-CoChR-GFP neurons in the APG are activated with patterned light (galvo-steered) while HR is recorded. **(O–Q)** Trial-averaged HR responses to APG activation at 5, 7, 12 dpf (gray, individual trials; red, mean; blue bar, stimulus). No effect at 5 dpf; increasing tachycardia at 7–12 dpf. Response is β-adrenergic-dependent (strongly reduced by propranolol; see Supp. Fig. 7). The shaded stimulation window marks optogenetic activation; concurrent heart-rate data were not acquired during this interval because stimulation interfered with cardiac imaging. **Analysis details.** Kernels estimated on the masked time base with causal fractional lags (0–4 s, 0.1 s step). Predictive neurons defined by Otsu threshold on per-neuron CV performance (with a 0.2 floor). GLM performance reported as CV R² (blocked folds). Clusters derived by k-means clustering in kernel PCA space (see Methods and Supp. Material, Clustering Kernels); motif means plotted with standard error (SE).

### Formation of Sympathetic Cardiac Circuits

In zebrafish, the sympathetic nervous system consists of a chain of paravertebral ganglia that extends from the visceral cavity to the tail (Fig. 4A). As in other vertebrates, these ganglia can be visualized using antibodies against tyrosine hydroxylase (TH), the rate-limiting enzyme in catecholamine biosynthesis [26]. To determine when the heart first receives sympathetic input, we performed TH immunostaining alongside Myl7 to label myocardial tissue. At 5 dpf, no TH-positive fibers were detected in the heart (Fig. 4C). By 7 dpf, TH-positive projections reached the sinoatrial plexus (SAP), indicating the onset of sympathetic cardiac innervation (Fig. 4C). These fibers originated from the most anterior sympathetic ganglia, which we refer to as the anterior paravertebral ganglia (APG) (Fig. 4C). Over development, the APG expand and adopt a distinctive C-shaped morphology (Fig. 4B). Despite this morphological maturation, TH-positive projections remain restricted to the SAP through 12 dpf, with no evidence of innervation to the atrioventricular plexus or ventricle.

To quantify the developmental dynamics and interindividual variability of sympathetic innervation, we assessed individual animals at 5, 7, and 12 dpf and recorded the presence or absence of TH-positive fibers at the heart. At 5 dpf, none of the nine animals examined showed sympathetic innervation. By 7 dpf, 6 of 13 animals exhibited TH-positive fibers at the SAP, indicating a variable onset of innervation across individuals. By 12 dpf, all 10 animals displayed consistent TH-positive projections to the SAP (Fig. 4D, E). These findings reveal a gradual and variable onset of sympathetic connectivity to the heart, with full anatomical incorporation of the APG circuit established by 12 dpf.

### Emergence of Control Motifs and Function in Sympathetic Cardiac Circuits

To investigate how the anterior paravertebral ganglia (APG) acquire the ability to regulate cardiac function, we expressed GCaMP6f under the TH1-Gal4 driver and we simultaneously imaged sympathetic neuron activity and heart rate dynamics in response to brief UV flashes (Fig. 5A). At 5 dpf, APG neurons respond to the UV stimulus with transient increases in activity (Fig. 5B), indicating that these neurons are already recruited by visual stress before direct sympathetic cardiac innervation is detectable. At 7 dpf, sympathetic neurons respond with apparent faster dynamics (Fig. 5C). By 12 dpf, as the ganglion increased in size, we observed emergent heterogeneity: newly appearing neurons decreased their activity following the same stimulus (Fig. 5D), hinting at the maturation of diverse sympathetic subtypes.

To further investigate the diversity of neural dynamics and their temporal relationship with heart rate, we trained a generalized linear model (GLM) to predict heart rate from the activity of all recorded APG neurons. This model estimates a temporal kernel for each neuron that captures how activity at different time lags contributes to prediction of heart rate. Pooling across animals revealed a striking diversity of kernel shapes (Fig. 5E). Unsupervised clustering of kernels identified six reproducible motifs (Fig. 5F, Supp. Fig. 6). Two motifs were biphasic with opposite polarity (C1, C4), resembling derivative filters that emphasize the effect of changes in neural activity. Two were monophasic bumps (C2, C5), consistent with proportional decoding of magnitude. The remaining two were monotonic ramps (C3, C6), consistent with integrator functions that accumulate activity over time. Together, these motifs map onto canonical motor control operations—derivative, proportional, and integrator—that may form the building blocks of cardiac control.

**Figure 6.**
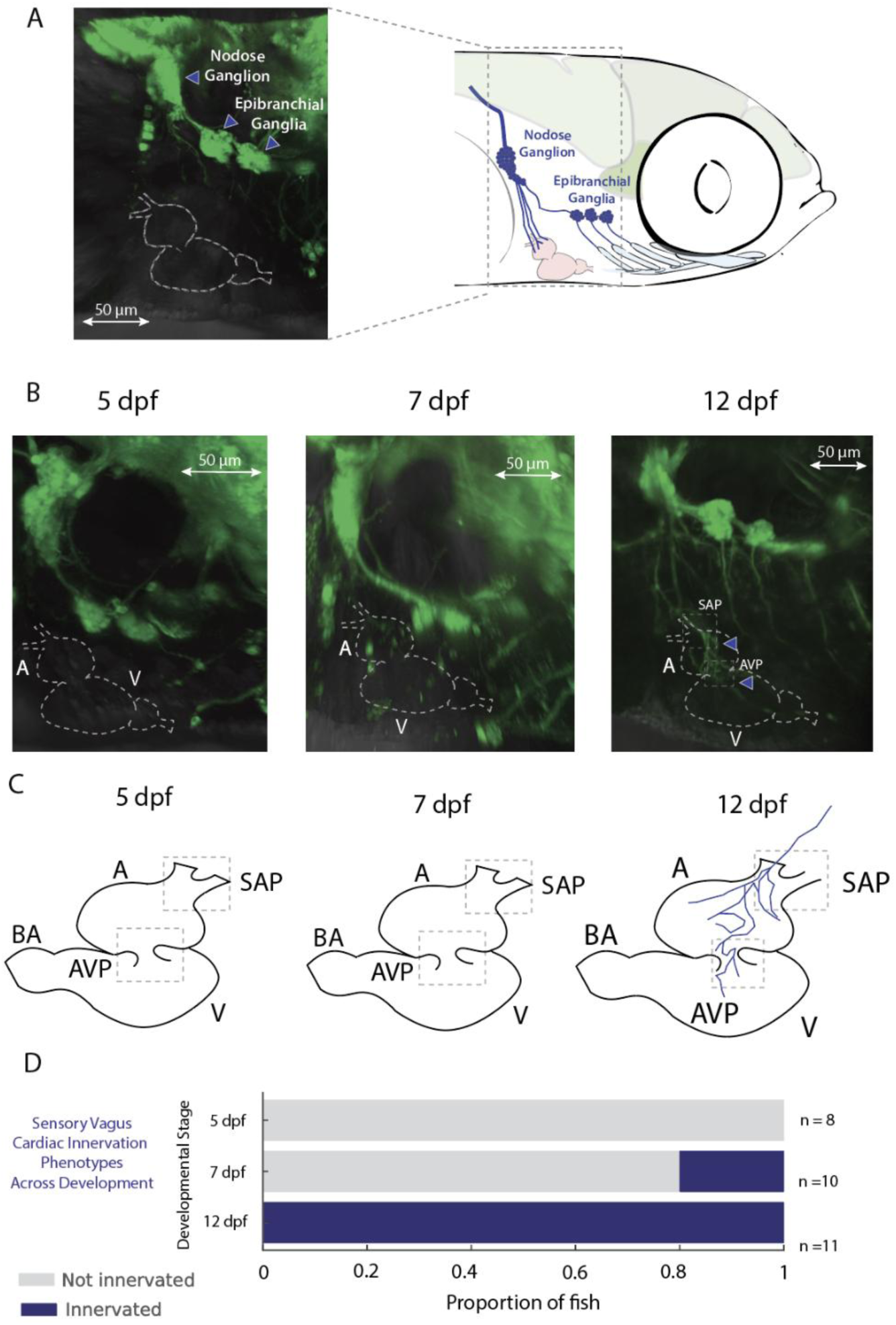
Developmental establishment of the heart-to-brain interoceptive circuit. (A) Vagal sensory anatomy. *Vglut->GFP* labels cranial sensory neurons; the nodose ganglion (posterior) and epibranchial ganglia (anterior) are indicated (left). Schematic (right) shows ganglia positions and sensory projections toward the heart. Scale bar, 50 µm. (B) *In vivo* imaging of vagal sensory projections to the heart at 5, 7, and 12 dpf. Dashed outlines mark atrium (A) and ventricle (V). 5 dpf: ganglia present but no cardiac sensory fibers. 7 dpf: sparse fibers in a subset of animals. 12 dpf: robust sensory projections reach the sinoatrial plexus (SAP) and atrioventricular plexus (AVP) (arrowheads). Scale bars, 50 µm. (C) Schematic summary by age: absence at 5 dpf, variable onset at 7 dpf, and established SAP/AVP innervation by 12 dpf. BA, bulbus arteriosus. (D) Population summary of sensory vagus cardiac innervation (presence/absence of heart-projecting fibers) across development. 5 dpf n=8: 0% innervated; 7 dpf n=10: 20% innervated; 12 dpf n=11: 100% innervated. **Acronyms:** A, atrium; V, ventricle; SAP, sinoatrial plexus; AVP, atrioventricular plexus; BA, bulbus arteriosus.

In our decoding model, neural activity is passed through its assigned kernel, summed across neurons, and then transformed by a static sigmoid nonlinearity to predict heart rate (Fig. 5G). Cross-validated performance improved significantly with age, with CV-R² increasing from 0.2 at 5 dpf to ∼0.5–0.6 at 12 dpf (Fig. 5H), indicating that APG neurons acquire predictive power as circuits mature. To illustrate the predictive power of the model even with modest CV-R^2^, we display the heart rate over time (in blue) and the heart rate predicted by the model (in orange) (Fig. 5I). Cluster composition also shifted across development: derivative motifs were absent early (except for one neuron) and emerged at 7 and 12 dpf (Fig. 5J), suggesting that the APG progressively establishes the computational repertoire needed for cardiac control.

To investigate whether specific neuron types are spatially organized in the ganglion, we plotted the imaged APGs coloring each neuron by cluster. We found that locations are variable per fish and per side, without a stereotypic location (Fig. 5K-M).

Last, to directly test the causal role of sympathetic neurons in modulating heart rate, we optogenetically activated APG neurons expressing CoChR under the TH1-Gal4 promoter. Using galvo mirrors, we delivered spatially targeted stimulation to the APG while monitoring heart rate immediately before and after stimulation; because of the APG’s location relative to the cardiac imaging field, heart rate could not be reliably measured during the stimulation period itself (Fig. 5N). At 5 dpf, stimulation failed to evoke any change in heart rate (Fig. 5O), consistent with the absence of sympathetic innervation. At 7 dpf, however, optogenetic activation induced modest heart rate increases (Fig. 5P), which became progressively larger by 12 dpf (Fig. 5Q). Quantification across animals confirmed this developmental emergence of APG-evoked tachycardia (Supp. Fig. 7).

**Figure 7.**
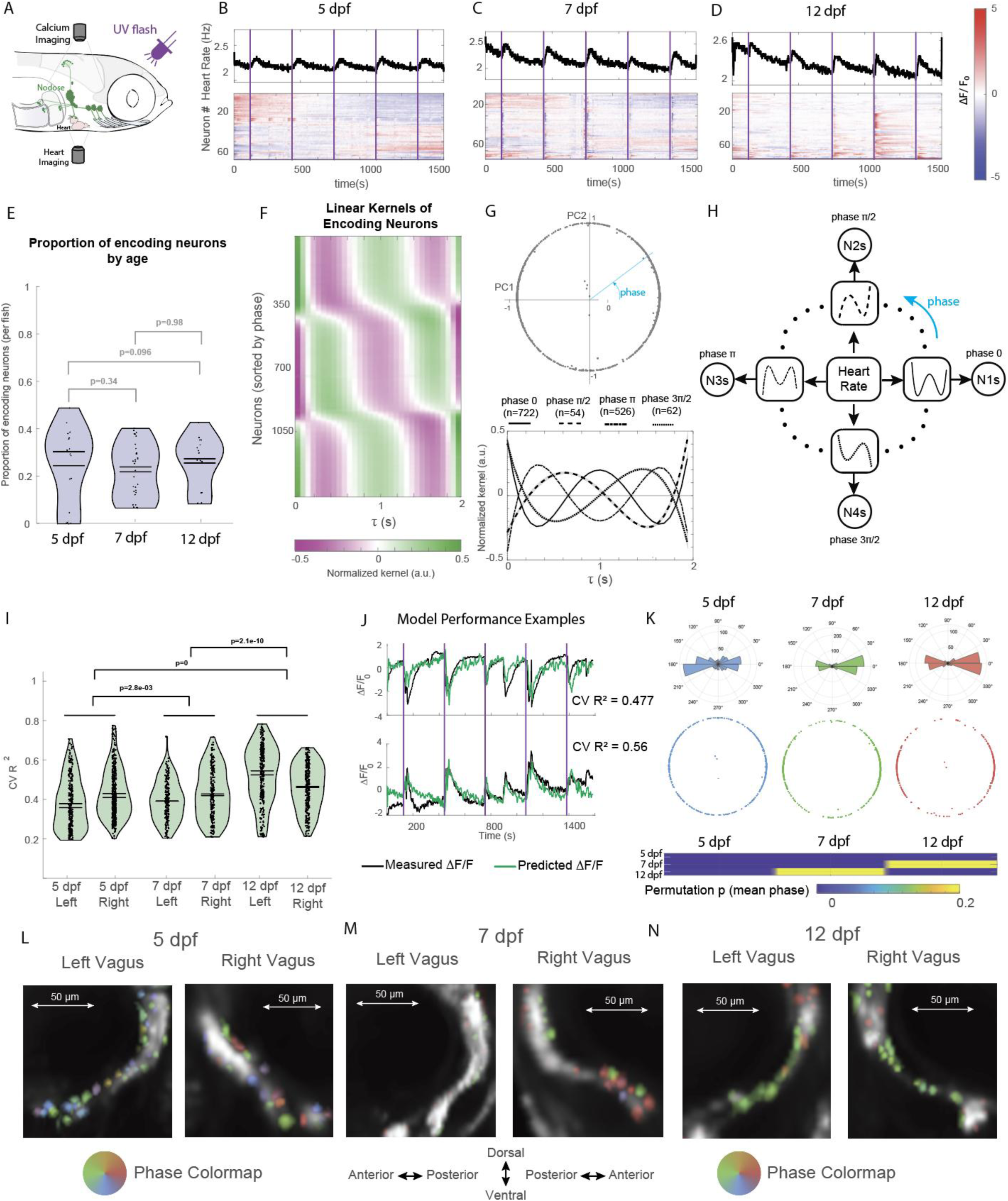
Cardiac encoding in vagal sensory neurons. **(A)** Experimental schematic. Calcium imaging of vagal sensory neurons (vglut2-Gal4>UAS-GCaMP6s) was performed simultaneously with optical heart-rate (HR) readout during UV-flash stimuli. **(B–D)** Representative recordings of HR (top, Hz) and vagal ganglion population activity (bottom, ΔF/F_0_ heatmaps; rows = neurons; columns = time) at 5, 7, and 12 dpf. Purple bars indicate stimuli. (E) Proportion of cardiac-encoding neurons across development. Encoding was defined as neurons with CV R² > 0.2 in GLM fits of HR→neural activity. Points are individual fish, violins show distributions, and p values are Wilcoxon rank-sum with BH-FDR correction. (F) Heatmap of normalized linear kernels from all encoding neurons, sorted by shape similarity. A single biphasic motif is expressed with systematic phase shifts. (G) Principal component analysis (PCA) of kernel shapes. Top: distribution of kernel phases in PC1–PC2 space lies along a circular trajectory. Bottom: canonical kernels at four representative phase bins (0, π/2, π, 3π/2). (H) Schematic representation of phase-shifted kernel ensembles. Neurons tile the canonical waveform across four quadrature-like groups. (I) Model performance across development. Violin plots show CV R² values (blocked 5-fold cross-validation) for left and right vagus at 5, 7, and 12 dpf. Horizontal bars denote medians; annotated p values are Wilcoxon rank- sum with BH-FDR correction. (J) Example neurons illustrating predictive power of the GLM. Black, measured ΔF/F0; green, predicted ΔF/F0. Top: neuron with phase π kernel decreases activity when HR rises. Bottom: neuron with phase 0 kernel increases activity when HR rises. (K) Radial histograms of kernel phase distributions at 5, 7, and 12 dpf. Early neurons are biased toward 0 and π; by 12 dpf distributions become bimodal, with encoders at opposite phases. Inset: permutation p-value heatmap showing significant differences in circular mean phase between 5 and 7 dpf and between 5 and 12 dpf. **(L–N)** Spatial distribution of encoding neurons colored by kernel phase in left and right nodose ganglia at 5 (L), 7 (M), and 12 (N) dpf. Neurons of different phases are intermixed within each ganglion and across both sides.

Importantly, the response was strongly reduced by propranolol, a β-adrenergic receptor antagonist (Supp. Fig. 7), confirming that sympathetic neurons modulate cardiac function via β-adrenergic signaling.

### Formation of the Heart-to-Brain Interoceptive Circuit

The somata of the sensory vagus are organized into four ganglia per side. The largest and most posterior is the nodose ganglion. It is located below the lateral line ganglion and innervates the heart and most of the viscera. The three most anterior ganglia are the epibranchial ganglia, which innervate each of the gill arches and are believed to track oxygen and other ventilatory parameters [28] (Fig. 6A). To visualize vagal ganglia, we expressed GFP under the control of the vglut promoter, which labels the cranial sensory neurons [29].

Central projections from the developing vagal sensory ganglia were already evident at 2, 3, and 4 dpf, extending into the hindbrain before peripheral organ innervation was complete (Supp. Fig. 8), consistent with previous anatomical descriptions of the larval zebrafish vagal sensory system [67]. To assess the developmental timing and variability of cardiac innervation by the sensory vagus nerve, we examined the presence of sensory axonal projections to the heart at 5, 7, and 12 dpf. At 5 dpf, all fish (8/8) had formed vagal sensory ganglia but lacked projections to the heart, indicating that ganglion specification precedes target innervation. By 7 dpf, only 2 of 10 animals exhibited cardiac sensory innervation, revealing substantial interindividual variability in the onset of sensory connectivity. By 12 dpf, all fish (11/11) showed robust vagal sensory projections to the heart, demonstrating that sensory innervation becomes fully established at this stage.

**Figure 8.**
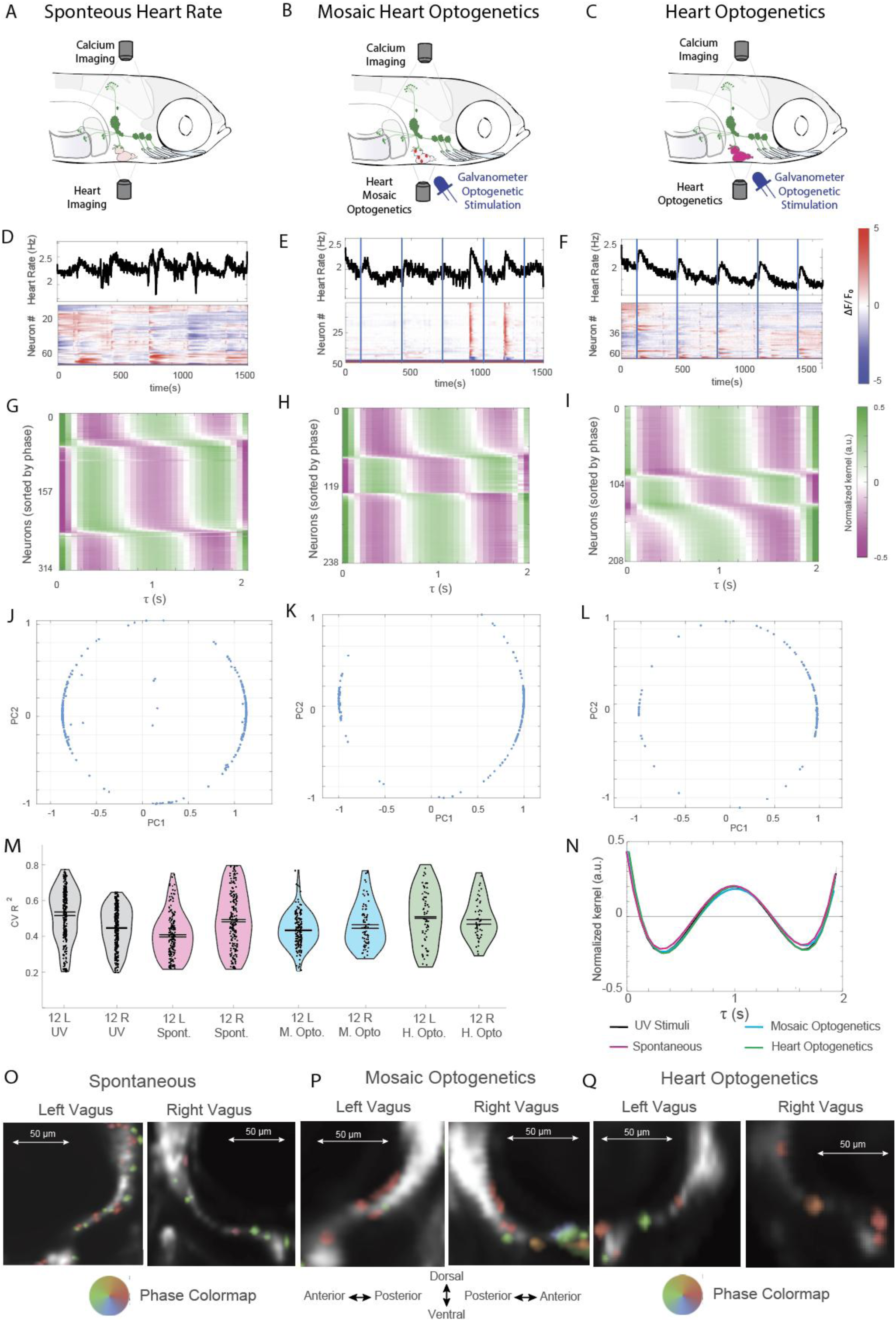
Stimulus-invariant encoding of heart rate in vagal sensory neurons. **(A–C)** Experimental paradigms for three sources of heart-rate modulation. (A) Spontaneous fluctuations in the absence of external perturbations. (B) Mosaic optogenetic activation of small subsets of cardiomyocytes (blue arrow, galvo-delivered stimulation). (C) Global optogenetic activation of the entire myocardium (blue arrow, galvo-delivered stimulation). Calcium imaging of vagal sensory neurons was performed simultaneously with optical readout of heart rate. **(D–F)** Representative recordings of heart rate (top, Hz) and vagal ganglion calcium activity (bottom, ΔF/F0 heatmaps; rows = neurons; columns = time) during (D) spontaneous fluctuations, (E) mosaic optogenetic perturbations, and (F) global optogenetic tachycardia. Blue bars mark stimulus onset in optogenetic conditions. **(G–I)** Heatmaps of causal kernels from GLMs trained to predict neural activity from HR dynamics under each condition. Kernels are aligned by phase, revealing the same biphasic motif expressed with systematic phase shifts, as observed under UV stimulation (Fig. 7F). **(J–L)** Principal component analysis of kernel shapes. Kernels distribute along a circular trajectory in PC1–PC2 space, consistent with phase-shifted versions of a single canonical filter. (M) Cross-validated model performance (CV R²) across conditions. Violin plots show distributions for left (L) and right (R) vagus at 12 dpf. Median encoding strength was comparable across conditions (0.4–0.55). (N) Canonical kernel shapes recovered across spontaneous, mosaic, and global optogenetic perturbations converge to the same biphasic temporal motif. UV stress condition from Fig. 7 is replotted for comparison. **(O–Q)** Anatomical maps of encoding neurons colored by kernel phase in left and right nodose ganglia under (O) spontaneous fluctuations, (P) mosaic optogenetic perturbations, and (Q) global heart optogenetic activation. Phase groups are intermixed within each ganglion and across both sides, with a bias toward ventral positions but no strict spatial segregation.

### Cardiac encoding in vagal sensory neurons

In order to investigate how vagal sensory neurons may encode heart rate dynamics, we expressed GCaMP6s under the control of the vglut2-Gal4 driver. We selected this driver because in zebrafish, all cranial sensory neurons, including vagal sensory neurons (VSNs), are glutamatergic [29]. We did not restrict our experiments to 12dpf – when vagal sensory innervation of the heart is established – because heart rate information could be available to VSNs from innervation of vasculature, gills, or other indirectly coupled structures. We stimulated fish with UV-flashes while recording calcium activity in the vagal ganglia concomitantly with heart rate (Fig. 7A). We found robust heart rate responses across development, from 5 to 12 dpf, and diverse neural dynamics, including subpopulations of VSNs that displayed temporally locked responses (Fig. 7B-D). Using generalized linear models (GLMs), we asked how well heart-rate fluctuations predict neural activity, assigning each neuron a causal temporal kernel. Neurons exceeding a cross-validated performance threshold (CV R² > 0.2) were classified as cardiac-encoding. The proportion of encoding neurons was comparable at 5, 7, and 12 dpf (Fig. 7E), indicating that subsets of VSNs track heart-rate trajectories even before direct sensory innervation of the heart is fully established.

Kernel heatmaps aligned by shape revealed that the diversity of response filters could be described as a single temporal motif expressed at systematically shifted phases (Fig. 7F). In other words, rather than multiple distinct kernel families, each encoding neuron appeared to express the same biphasic kernel advanced or delayed in time. Principal-component analysis confirmed this interpretation: the first two PCs accounted for nearly all variance (Supp. Fig. 9) and arranged the kernels along a circular trajectory in phase space (Fig. 7G). This geometry is consistent with a basis set of sinusoidal quadrature components, where neurons are distributed around the circle such that each one samples a different phase of the same underlying waveform (Fig. 7H).

**Figure 9.**
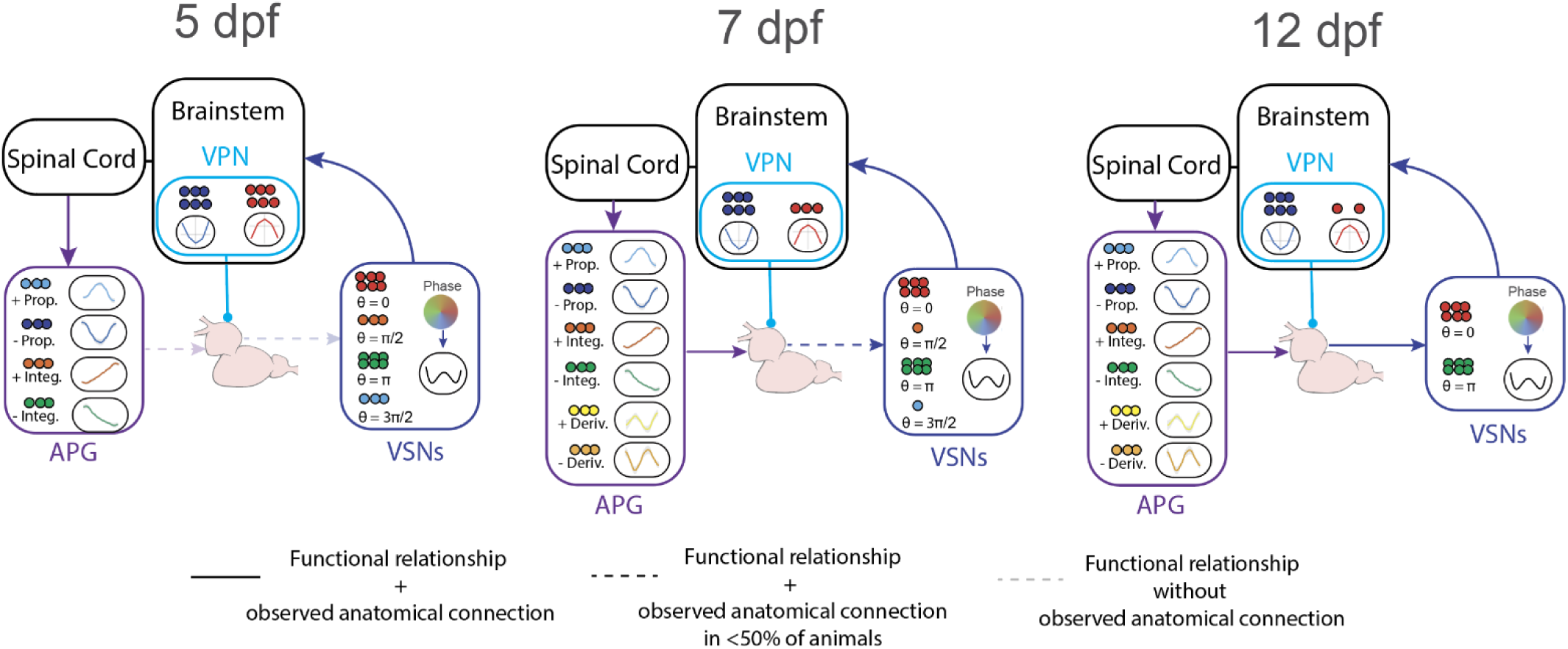
Emergence of functional heart–brain circuits. Graphical summary of the staged assembly of cardiac autonomic circuits across larval development. Arrows indicate functional relationships identified by encoding, decoding, and/or perturbation experiments. Solid arrows indicate functional relationships for which anatomical connections were observed at the corresponding developmental stage. Dark dashed arrows indicate functional relationships for which anatomical connections were observed in fewer than half of the animals examined. Light dashed arrows indicate functional relationships for which a corresponding anatomical connection was not observed in this study at that stage. Kernel schematics summarize the dominant temporal motifs identified in each neuronal population. At 5 dpf, descending vagal motor pathways provide early inhibitory control of heart rate. By 7 dpf, neurons in the anterior paravertebral ganglia (APG) provide sympathetic modulation and exhibit proportional, integrative, and derivative-like motifs. By 12 dpf, vagal sensory neurons (VSNs) innervate the heart and encode heart-rate dynamics through a phase-shifted population code. The heart is intrinsically autorhythmic; the neural pathways shown here modulate an ongoing rhythm, and endocrine, paracrine, metabolic, and other non-neuronal influences are not depicted. VPN, functionally defined vagal premotor locus neurons.

The quality of this representation improved across development. Model performance increased, with cross-validated (CV) R² values rising from ∼0.3 at 5 dpf to >0.5 by 12 dpf (Fig. 7I), indicating that HR became a progressively stronger predictor of neural activity. To illustrate the predictive power of the model even with CV- R^2^ values near the median, we display the ΔF/F_0_ of a neuron with phase π (top) and a neuron with phase 0 (bottom) over time (in black) and the ΔF/F_0_ predicted by the model (in green) (Fig. 7J). In both cases the signal is tracked accurately, with every large fluctuation in measured ΔF/F_0_ matched by a large fluctuation in the predicted ΔF/F_0_. This is also important to illustrate that neurons with phase π decrease activity when heart rate increases following the dynamics of the canonical kernel, and neurons with phase 0 increase activity when heart rate increases, also following the dynamics established by the canonical kernel.

The distribution of kernel phases also changes systematically during development: at 5 dpf, neurons did not occupy phases uniformly, but instead showed a bias toward 0 and π (Fig. 7K, blue radial histogram). By 7 dpf, these biases strengthened (Fig. 7K, green radial histogram), and by 12 dpf neurons clustered into two modes of representation: a positive biphasic kernel (at phase 0) and a negative biphasic kernel (at phase π) (Fig. 7K red radial histogram). We confirmed an age-dependent shift in kernel phase using pairwise permutation tests of circular mean phase, which showed significant differences between 5 dpf and both 7 and 12 dpf (Fig. 7K inset heatmap), consistent with the developmental reorganization of phase representation.

Anatomical mapping showed no spatial segregation of phase groups: neurons from different ensembles were intermixed within vagal ganglia and across both sides (Fig. 7L–N). Thus, the phase structure of the code is functional rather than anatomical, arising from temporal offsets in kernel phase rather than physical clustering.

Together, these findings demonstrate that vagal sensory neurons implement a distributed, phase-shifted population code for heart rate. By expressing a single canonical kernel at different phase offsets, the ensemble provides a robust and redundant representation of cardiac dynamics, well before direct cardiac sensory innervation is fully stabilized, and, as the fish matures, the encoding shifts to a bimodal representation with kernels of opposite phases.

### Stimulus-invariant heart rate encoding in vagal sensory neurons

The observation that a single kernel shape, expressed at shifted phases, accounts for heart rate encoding across fish and developmental stages suggests that this is a core computational property of vagal afferents. To test this hypothesis, we asked whether the same phase-shifted kernel emerges under distinct modes of heart-rate modulation: (1) spontaneous fluctuations, which occur without environmental perturbations (Fig. 8A); (2) fluctuations elicited by mosaic optogenetic activation of small subsets of cardiomyocytes (Fig. 8B); and (3) acute tachycardia triggered by brief optogenetic activation of the entire myocardium (Fig. 8C). Spontaneous fluctuations were chosen because, in contrast to fight-or-flight responses, they occur while the external sensory environment is constant and therefore span a broad range of amplitudes, durations, and temporal profiles (Fig. 8D). Optogenetic manipulations were included because they perturb the heart directly and independently of external stimuli. For these experiments, we generated transgenic zebrafish expressing the light-gated cation channel CoChR under the cmlc2 promoter, which labels cardiomyocytes [30]. Mosaic activation produced local perturbations that did not evoke immediate tachycardia but instead generated delayed fluctuations, sometimes tens of seconds after stimulation (Fig. 8E). In contrast, global heart activation produced immediate and robust tachycardia (Fig. 8F). Together, these three conditions provided diverse trajectories of heart rate and neural activity (Fig. 8D–F) without altering the external environment, allowing us to probe encoding mechanisms under controlled and distinct perturbations.

We next trained GLMs, as in Fig. 7, to identify causal kernels that predict neural activity from heart-rate dynamics. Across all conditions, kernel heatmaps revealed the same smooth green↔magenta gradient (Fig. 8G–I) seen previously under UV stress (Fig. 7F), indicating that most encoding neurons are well described by a single canonical biphasic filter, differing primarily in phase (latency/polarity). PCA of kernel shapes confirmed this structure: in all three conditions, kernels distributed along a circular trajectory in PC space, with neurons clustering near phase 0 and π (Fig. 8J–L), as observed in 12 dpf fish during UV stimulation (Fig. 7K). Encoding strength was similar across conditions, with median CV R² values in the 0.4–0.55 range (Fig. 8M). Moreover, the canonical kernel identified in each case converged to the same shape (Fig. 8N).

As in the neural recordings of responses to UV flashes, anatomical mapping showed no spatial segregation of phase groups: neurons from different ensembles were intermixed within vagal ganglia and across both sides (Fig. 8O–Q). Across fish, we observe a higher likelihood of neurons being located on the ventral side of the nodose ganglion, however, there are also neurons in more dorsal parts of the ganglia (Fig. 7L-N, Fig. 8O- Q).

These results demonstrate that vagal sensory neurons implement a stimulus-invariant, phase-shifted (latency/polarity) population code for heart rate. Whether fluctuations arise from environmental stressors, spontaneously, from localized perturbations, or from optogenetically-induced tachycardia, the dynamics of VSN encoding are well described by a linear time-invariant filter set, with phase (latency/polarity) offset as the primary degree of freedom.

## Discussion

Here, we have defined the developmental sequence by which heart–brain functional circuits emerge and mature in a vertebrate. Using larval zebrafish, we combined longitudinal calcium imaging, optogenetics, quantitative optical cardiophysiology, and computational modeling to dissect the limbs of the autonomic circuit. We show that descending vagal motor projections provide the first detected neural efferent pathway for cardiac modulation before intracardiac neurons appear; that a functionally defined cholinergic hindbrain locus engages early cardioinhibitory control; that sympathetic inputs from the anterior paravertebral ganglia add modulatory capacity and their dynamics display proportional, integrative, and derivative-like control motifs; and that vagal sensory neurons encode heart rate with a canonical biphasic kernel expressed at distinct phase offsets, yielding a robust, stimulus-invariant population code. Together, these findings reveal how neural regulation of the heart is assembled in stages and how each stage contributes a distinct computational role.

### Early brain-to-heart communication via the motor vagus

Our findings identify the descending vagal pathway as the first detected neural efferent pathway capable of modulating heart rate in zebrafish (Fig. 1 and 2). By 5 dpf, cholinergic fibers reach the sinoatrial plexus, while sympathetic cardiac innervation and intracardiac neurons are not yet detectable. Targeted activation of the cholinergic hindbrain locus described in Fig. 3 elicits bradycardia once cardiac innervation is present, supporting engagement of descending vagal cardiac inhibition. This stage is particularly striking because it separates the earliest onset of neural control from the later incorporation of intracardiac ganglia. In mammals, parasympathetic ganglia embedded in the cardiac plexus are thought to be essential for modulation of pacemaker activity [31, 32], but our results suggest that brain-to-heart projections alone can transiently carry this function. Such a configuration may provide a rapid developmental solution for ensuring that cardiac activity is already under neural control during the critical window when larvae first engage with the external environment.

### A cholinergic premotor locus for descending control

In parallel with the onset of vagal motor projections, we identified a small medial cluster of hindbrain neurons whose activity reliably predicts cardiac inhibition. Encoding models revealed a canonical kernel with negative polarity, and targeted optogenetic activation of this locus elicited bradycardia in animals as early as 7 dpf (Fig. 3S). These data indicate that a functionally defined premotor locus in the hindbrain identifies a reproducible functional node for descending cardiac control. We use the term premotor because the neurons were identified by their functional relationship to heart-rate dynamics and by targeted perturbation, rather than by single-cell anatomical reconstruction. Thus, this locus may include motor vagus neurons, neurons upstream of motor vagus neurons, or both. Regardless of this cellular composition, its cholinergic identity, medial hindbrain location, predictive dynamics, and ability to elicit bradycardia support the conclusion that it engages the descending vagal pathway for cardiac inhibition. The cluster-restricted decoding analysis further shows that heart-rate-related predictive information is present in both opposing kernel classes (Supp. Fig. 4).

### Sympathetic innervation and emergence of control motifs

The second stage of circuit assembly involves the arrival of sympathetic projections from the anterior paravertebral ganglia (Fig. 4). Anatomical innervation emerges and optogenetic activation shows that sympathetic neurons acquire the capacity to accelerate the heart at 7 dpf. At 5 dpf, however, APG neurons already respond to visual stress and covary with heart-rate dynamics, despite the absence of TH-positive cardiac innervation and APG-evoked tachycardia. We therefore interpret this early relationship as coordinated recruitment of the APG and the cardiac response within a broader stress-related physiological state rather than as direct sympathetic control of the heart. Endocrine stress pathways, including the hypothalamic–pituitary– interrenal axis, may also contribute to this coordinated response at 5 dpf, although we did not directly test their involvement. Encoding analyses of APG activity uncover kernels with proportional, integrative, and derivative-like temporal motifs. This computational repertoire suggests that the sympathetic system contributes not only to the magnitude but also to the dynamics of cardiac responses. Such motifs may allow the system to respond proportionally to changes in heart rate, integrate sustained deviations, or accentuate rapid fluctuations, thereby shaping the gain and stability of autonomic control. Mammalian baroreflex studies indicate modulation of gain and sensitivity that is consistent with proportional control, and dynamic changes in reflex latency under different physiological conditions, suggesting that components of the control motifs we observe may be conserved [33, 34]. Our findings are the first evidence for heart rate integral- or derivative-like temporal dynamics in sympathetic neurons.

### Development of the intracardiac nervous system

Although our primary focus was on the staged emergence of autonomic control, our results also provide new insight into the developmental progression of the intracardiac nervous system (ICNS). By tracking phox2bb+ intracardiac neurons alongside functional assays of heart rate control, we found that cardiac innervation is first detectable at 5 dpf, but intracardiac neurons are absent at this stage. The earliest intracardiac neuron appears by 7 dpf, and by 12 dpf multiple clusters are evident, including a reproducible population in the dorsal–posterior atrium that we designate the intracardiac ganglion. Thus, the arrival of ICNs lags behind the onset of functional vagal efferent control, which is already functional by 5 dpf. This indicates that the initial establishment of parasympathetic influence over heart rate occurs independently of intracardiac ganglia, with these neurons contributing only after the basic circuit is in place.

In mammals, intrinsic cardiac ganglia are thought to be essential for modulation of pacemaker activity and for shaping the spatiotemporal dynamics of cholinergic input to the sinoatrial and atrioventricular nodes [31, 32]. Our findings suggest that in zebrafish, these neurons are dispensable for the earliest phases of functional control. They likely refine or diversify cardiac modulation as they integrate into the circuit. This sequence may reflect a developmental principle: early reliance on extrinsic parasympathetic fibers to ensure inhibitory capacity, followed by later recruitment of intracardiac ganglia for fine-tuned control.

### Sensory feedback and the establishment of heart rate encoding

The final stage in the developmental sequence is the engagement of vagal sensory neurons (VSNs). By 12 dpf, sensory axons reliably reach the sinoatrial and atrioventricular plexuses, but encoding analyses revealed that VSNs can track heart-rate fluctuations even at earlier ages, likely through indirect coupling via vasculature, gill innervation or innervation of other tissues with heart rate-coupled dynamics (Fig. 6-8). Across spontaneous fluctuations, responses to visual threat, local optogenetic perturbations, and optogenetically-induced tachycardia, a single biphasic kernel accounted for neural responses. Neurons differed only in phase, tiling the kernel at systematic offsets. This phase-shifted organization sharpened with development, evolving into a bimodal distribution centered at opposite polarities. Such a population code allows continuous coverage of heart-rate dynamics while remaining robust to the source of modulation. The emergence of this stimulus-invariant representation demonstrates that VSNs contribute a stable and generalizable code for cardiac interoception.

Prior studies have described heartbeat-locked activity and cardiovascular convergence in central brainstem neurons, including neurons in the nucleus tractus solitarius [68,69]. These responses reflect neural activity aligned to individual cardiac cycles. In contrast, the encoding described here occurs on the slower timescale of heart-rate dynamics: VSN activity tracks changes in beat frequency over seconds across spontaneous fluctuations, visual threat responses, and optogenetically induced cardiac perturbations. We term this computational property chronoreception, by analogy to chemoreception and baroreception, and consistent with the use of “chronotropic” to refer to changes in heart rate [35]. Hemodynamically, direct information about heart rate is potentially useful because, together with stroke volume, heart rate contributes to cardiac output.

Regardless of whether heart-rate dynamics are detected at the heart or through other mechanically coupled structures, these encoders are present across conditions and provide afferent signals to the brain.

### Integration of efferent and afferent development into a staged program

Together, our findings reveal a staged program for the assembly of heart–brain circuits (Fig. 9). The first functional stage is established by descending cholinergic vagal innervation, which enables neural modulation of heart rate before intracardiac neurons or sympathetic cardiac projections are present. A medial cholinergic hindbrain locus engages this early inhibitory control. Intracardiac neurons are subsequently incorporated into the circuit, while sympathetic innervation reaches the heart and APG neurons acquire the ability to accelerate heart rate. As this sympathetic pathway matures, cardiac responses increase in amplitude and speed, and APG activity develops a broader repertoire of proportional, integral, and derivative-like relationships with heart- rate dynamics. The afferent limb emerges later; however, vagal sensory neurons represent heart-rate dynamics across spontaneous and experimentally induced perturbations even before stable cardiac innervation is established, and subsequently form robust projections to the cardiac plexuses. Together, these stages transform an initially minimal descending pathway into a feedback control system capable of regulating cardiac function and representing its consequences.

### Limitations and open questions

Several important questions remain about how this developing control system is connected and how cardiac state is sensed. The functional relationships identified through encoding, decoding, and perturbation do not by themselves establish direct synaptic connectivity, and the specific postsynaptic targets of vagal sensory projections within the hindbrain still need to be identified. Likewise, the peripheral structures through which VSNs detect heart-rate-related dynamics have not yet been resolved. Although sensory axons reliably reach the cardiac plexuses by 12 dpf, VSNs represent heart-rate fluctuations at earlier stages, suggesting that these signals may also arise through vascular, gill, or other mechanically coupled tissues. Connectomic approaches, building on the ultrastructural analysis used here for the developing intracardiac circuit, should help resolve how these afferent signals are integrated with vagal and sympathetic efferent pathways.

### Comparisons to mammalian autonomic development

The sequence we describe parallels aspects of autonomic development in mammals and creates new opportunities for studying how these circuits become functional. In rodents and humans, autonomic innervation of the heart arises prenatally, making it difficult to observe the onset of function *in vivo* [7]. Fetal studies suggest that vagal tone emerges early and that sympathetic influences strengthen later in gestation [36].

Sensory encoding of blood pressure has been inferred from nodose baroreceptor recordings in mammals, which respond to beat-to-beat pressure oscillations [37, 38], but direct demonstration of heart rate dynamic encoding has been lacking. Our results provide the first continuous, cell-resolved account of how HR is encoded in a vertebrate vagal sensory ganglion.

### Implications for interoception and autonomic disorders

Recent studies have shown that interoceptive signals from the heart influence emotion, cognition [39], and behavior [40], and disruptions in cardiac sensory coding have been implicated in conditions ranging from anxiety to dysautonomia [41,42]. Cardiac interoception may also be relevant to cardiovascular diseases in which cardiac dynamics are altered acutely or chronically, including arrhythmias, heart failure, and myocardial infarction. In these conditions, changes in rhythm, contractility, perfusion, and autonomic tone alter the physiological signals available to cardiac afferents and may shape compensatory reflexes, symptom perception, and behavioral state. This sensory limb is clinically underexplored [42]: current cardiovascular therapies largely target cardiac tissue or efferent mechanisms that alter heart rate, contractility, or vascular tone, whereas cardiac sensory and interoceptive pathways are rarely targeted directly. Our identification of a cell-resolved vagal sensory code for heart-rate dynamics provides an entry point for testing how afferent representations are altered by cardiac dysfunction and whether modifying heart-to-brain signaling can influence maladaptive autonomic responses. By establishing zebrafish as a model in which efferent and afferent components can be interrogated optically, this work enables direct analysis of how genetic, developmental, physiological, or injury-induced perturbations reshape interoceptive coding.

## Conclusion

By resolving heart–brain circuits as they assemble, this study reveals how successive autonomic components add distinct control and sensory functions to a developing physiological system. Development provides a natural experimental decomposition of the circuit, allowing the contribution of individual components to be identified before they become fully interconnected in the mature animal. More broadly, our work establishes larval zebrafish as a vertebrate model for brain–body interactions in which neural activity, autonomic control, organ physiology, sensory feedback, and behavior can be observed and manipulated together in an intact animal at cellular resolution. This framework opens brain–body physiology to the experimental accessibility that has transformed other areas of systems neuroscience, creating opportunities to uncover general principles of interoception and neural control of internal organs.

## Acknowledgments

We thank members of the Hernandez-Nunez Lab for fruitful discussions and suggestions, in particular Shulin Zhang, Aliya Ablitip, Renzo Mendoza, Ting Ting Yan, and Sofia Melnychuck. We also want to thank members of the Engert and Fishman Labs: Vickie Wang, Yasuko Isoe, Kristian Herrera, Soma Singareddy, and Erin Song for early contributions and for useful discussions and feedback on this project. We would like to thank Richard Schalek and Juan-Carlos Tapia for their help with the serial section electron microscopy sample preparation and imaging. We thank the Harvard Center for Biological Imaging (RRID:SCR_018673) for infrastructure and support.

Luis Hernandez-Nunez received funding from the Harvard Mind Brain and Behavior Fellowship and Young Investigator Award, by the Life Sciences Research Foundation/Additional Ventures Fellowship, by the Warren Alpert Distinguished Scholar Award, by the Branco Weiss Fellowship, by the Burroughs Wellcome CASI grant, and by the National Institutes of Health (R34NS138096). Florian Engert received funding from the National Institutes of Health (U19NS104653 and R01NS124017), and the Simons Foundation (SCGB 542973 and NC-GB-CULM-00003241-02). Mark Fishman received funding from the National Institutes of Health (U19NS104653 and R01NS124017).

## Author Contributions

L.H.-N., F. E., and M. C. F. designed the study, interpreted the results, and wrote the manuscript with feedback from all authors. L.H.-N. developed the instruments and experimental techniques; wrote the software for data analysis; analyzed data; performed anatomical, behavioral, physiological, functional imaging, and optogenetic experiments; and built the encoding and decoding models. J.A. performed physiological, functional imaging, and optogenetic experiments related to the Sensory Vagus System (Fig. 7 and 8) with guidance from L.H.-N. S.S. performed physiological, functional imaging, and optogenetic experiments related to the Motor Vagus System (Fig. 2 and 3) with guidance from L.H.-N. A. M. performed physiological, functional imaging, and optogenetic experiments related to the Sympathetic System (Fig. 4, 5) with guidance from L.H.-N. A.K. conducted anatomical experiments of the Sensory Vagus Nerve that were the initial step for the results of Fig. 6 with guidance from L.H.-N. J. B.-W. contributed the electron microscopy reconstruction of the sinoatrial plexus of 7 dpf fish (Fig. 2G-I). V.R. and M.A. contributed early results on optogenetic activation of the motor vagus nerve, which were the initial steps toward Fig. 3R and S. A. Z.-S. generated the transgenic fish used for Fig. 8.

## Materials and Methods

### 1. Key Resource Table

| Reagent or Resource | Source | Identifier |
| --- | --- | --- |
| **Antibodies and chemicals** |  |  |
| Antigen: Acetylated alpha-tubulin. Host: mouse | Sigma-Aldrich. T6793 | link |
| Antigen: Tyrosine hydroxylase Host: mouse | Immonostar. 22941 | link |
| Antigen: Myl7 Host: rabbit | Sigma-Aldrich. HPA013331 | link |
| Propranolol hydrochloride powder >99%TLC | Sigma-Aldrich. P0884 | link |
| Atropine | Supelco. A-046 | link |
| **Zebrafish strains** |  |  |
| Tg(chata:Gal4VP16)mpn202 | Förster et al., 2017 | [46] |
| Tg2(phox2bb: EGFP) | Nechiporuk et al. 2007 | [47] |
| Tg(UAS:GCaMP6s) | Muto and Kawakami. 2011 | [48] |
| Tg(vglut2-Gal4, UAS:GCaMP6s) | Filippi et al. 2014 | [49] |
| Tg(ZK10205-TH1:Gal4-KI) | Kim et al. 2021 | [50] |
| Tg(cmlc2:GFP) | Huang et al. 2003 | [51] |
| Tg(UAS:GCaMP6f) | Janelia Farm Research Campus (JFRC) | [52] |
| Tg( UAS:CoChR-eGFP) | Janelia Farm Research Campus (JFRC) | [52] |
| Tg(vglut2-Gal4, UAS:GCaMP6s, myl7-UAS-CoChR-tdTomato) | This study |  |
| **Software** |  |  |
| Python 3.11 | Python | link |
| Matlab | Mathworks | link |
| ImageJ | NIH | link |
| CalmAn | Giovanucci et al (2019) | [53] |
| Zen Microscopy Software | Zeiss | link |

### 2. Table with the age and number of animals per experiment

Here we report the number of animals used for results that do not explicitly have the number of animals in the main figure.

| Figure | Age | Number of animals |
| --- | --- | --- |
| Figure 1C | 4 dpf | 15 |
| Figure 1C | 5dpf | 15 |
| Figure 1C | 7dpf | 20 |
| Figure 1C | 12dpf | 18 |
| Figure 1D | 4 dpf | 14 |
| Figure 1D | 5dpf | 16 |
| Figure 1D | 7dpf | 18 |
| Figure 1D | 12dpf | 16 |
| Supp. Figure 1C | 4 dpf | 20 |
| Supp. Figure 1D | 5dpf | 20 |
| Supp. Figure 1E | 7dpf | 20 |
| Supp. Figure 1F | 12dpf | 20 |
| Figure 3I | 5dpf | 12 |
| Figure 3I | 7dpf | 10 |
| Figure 3I | 12dpf | 9 |
| Figure 3R | 4dpf | 6 |
| Figure 3S | 7dpf | 6 |
| Figure 5H | 5dpf | 8 |
| Figure 5H | 7dpf | 11 |
| Figure 5H | 12dpf | 10 |
| Figure 5O | 5dpf | 24 |
| Figure 5P | 7dpf | 16 |
| Figure 5Q | 12dpf | 30 |
| Supp. Figure 7 | 5dpf Prop | 16 |
| Supp. Figure 7 | 5dpf Control | 16 |
| Supp. Figure 7 | 9dpf Prop | 11 |
| Supp. Figure 7 | 9dpf Control | 17 |
| Figure 7E | 5 dpf | 17 |
| Figure 7E | 7 dpf | 25 |
| Figure 7E | 12 dpf | 18 |
| Figure 8J | 12 dpf | 12 |
| Figure 8K | 12 dpf | 10 |
| Figure 8L | 12 dpf | 10 |

## Experimental model and subject details

### Zebrafish housing and maintenance

For all experiments, we raised small groups of 20–30 larvae in filtered fish water (exchanged daily) in Petri dishes (diameter: 9 cm) at a constant 28 °C. After 5 dpf, larval fish were fed with paramecia once per day, after 9dpf twice per day. We performed experiments with larvae at ages 2-12 dpf for developmental experiments. All animals were kept in accordance with The Harvard Institutional Animal Care and Use Committee (IACUC) [54]. The sex of the fish tested was unknown because it is only distinguishable after 20 dpf [55].

### Transgenesis

Transgenic zebrafish larvae were in casper or nacre background [56]. The fish *(UAS:GCaMP6f)^jf46^* and *(UAS:CoChR-eGFP)^jf44^* are described in Mu et al.[52]. The experiments that did not involve calcium imaging were conducted with *Nacre* fish. Anatomical observations were conducted using the progeny of *Tg(chata:Gal4VP16, UAS:GFP)^mpn202^* and *Tg2(phox2bb: eGFP)* for Figures 1 and 2 and the respective supplementary figures. For anatomical imaging of central vagal sensory projections (Supp. Fig. 8), we used Tg(vglut2a-Gal4; UAS-GFP) larvae.

For calcium imaging, we used transgenic zebrafish lines expressing:

**•** Cytosolic GCaMP6s in glutamatergic neurons, Tg(vglut2-Gal4, UAS:GCaMP6s) (Filippi et al.[49]) (Fig. 7,8)
**•** Cytosolic GCaMP6s in cholinergic neurons, Tg(chata:Gal4VP16, UAS:GCaMP6s)^mpn202^ (Förster et al.[46]) (Fig. 3)
**•** Cytosolic GCaMP6f in cells that express Tyrosine Hydroxylase, Tg(ZK10205-TH1:Gal4-KI, UAS:GCaMP6f) (Kim et al. [50] and Mu et al. [52]) (Fig. 5)

For optogenetic experiments, we used transgenic zebrafish lines expressing:

- CoChR in Tyrosine Hydroxylase cells Tg(ZK10205-TH1:Gal4-KI, UAS:CoChR-eGFP) (JFRC) (Fig. 5)
- CoChR in cholinergic neurons, Tg(chata:Gal4VP16, UAS:CoChR-tdTomato) (Fig. 3)
- CoChR in cardiomyocytes Tg(vglut2-Gal4, UAS:GCaMP6s, myl7-UAS-CoChR-tdTomato) (this study) (Fig. 8)
- For mosaic optogenetic experiments, we used the injected embryos (f0) from vglut2-Gal4, UAS:GCaMP6s, myl7-UAS-CoChR-tdTomato.

## Experimental Methods

### Immunohistochemistry

For myl7 and acetylated-alpha tubulin staining, animals were fixed in 4% PFA (Sigma-Aldrich) in PBS (Gibco, Grand Island, NY, USA). Primary antibodies were diluted in 3% bovine serum albumin in PBS and incubated at 4°C overnight. Secondary antibodies were diluted in 0.1% Triton in PBS and incubated for 2 h at room temperature. Alexa Fluor 488 was used for acetylated-alpha tubulin or Tyrosine Hydroxylase and Alexa Fluor 546 for myl7.

### Drug treatment

Propranolol treatments for Supp. Fig. 7 were systemic, with fish being placed in filtered fish water solutions of propranolol for 2 hours before measuring their responses to optogenetic stimulation of the APG. We used propranolol from Sigma-Aldrich (P0884). The concentration used was 20 µM of Propranolol (Supp. Fig. 7). One hour before the hemodynamic measurements, fish were partially immobilized using 2% agarose gel (Invitrogen, 16520). After 5 minutes of agarose solidifying, the tails of fish were released using a feather micro ophthalmic scalpel (Fisher Scientific link). Partially restrained fish were also in filtered fish water solutions of propranolol after embedding and during the heart rate measurements. The drug was washed out and the fish were tested for the control measurements 3-5 hours after the initial measurement. The concentration of propranolol was selected based on the reported concentrations used in [57]. At 9 dpf, 17 fish were exposed to propranolol. Because recordings during drug exposure were time-limited, APG-evoked cardiac responses were obtained from 11 animals during propranolol treatment. Following washout, responses were obtained from all 17 animals.

### Anatomical imaging

#### For fixated samples

Fixated samples were mounted on bottom glass Petri dishes (Thermo Fisher Scientific 150680) using 1.2% agarose and positioned using plastic tips. Mounted samples were then imaged using a Zeiss LSM 900 inverted scanning confocal microscope. Images were later processed in Zen (Zeiss microscopy software) and ImageJ to reconstruct the volumes.

#### For live animals

Live zebrafish were embedded in 1.8% low melting point agarose between 1 and 2 hours before imaging. They were positioned laterally using plastic tips on a bottom glass Petri dish. The fish were then imaged using a Zeiss LSM 900 inverted scanning confocal microscope. The step size and speed of recording were not set to the optimal confocal setting as we did for fixated samples. Instead, we selected a speed that allowed visualization of neurites that move with the heart as it beats (Supp. Vids. 1, 2, 3). Before imaging, the most fluorescent fish were selected by screening them using an Olympus MVX10 epifluorescence microscope. For fish that were imaged on both sides, fish were recovered after the first round of imaging using a micro ophthalmic scalpel to release the fish and then embedded on the other side after 1-2 hours.

### Serial section electron microscopy

The following protocol was specifically modified from Tapia et al.[58] to enhance extra-cellular space preservation, which improves synapse detection [59]. Unless otherwise noted, all steps were performed at room temperature (RT). A 7 dpf larva was anesthetized and embedded in agarose. The water was replaced with a dissection solution (64 mM NaCl, 2.9 mM KCl, 10 mM HEPES, 10 mM glucose, 164 mM sucrose, 1.2 mM MgCl2, 2.1 mM CaCl2, pH 7.5) supplemented with 0.02% tricaine3. Small slits were cut in the agarose to expose the eyes, and bilateral enucleations were performed to enhance ultrastructural preservation of the extra-cellular space and improve heavy metal staining. A custom-made hook was carefully inserted behind the eyes to minimize brain damage. The larva was then immediately transferred to a cold (4°C) fixation solution composed of 2.5% glutaraldehyde in 0.1 M cacodylate buffer (pH 7.4) supplemented with 4.0% mannitol. The cacodylate buffer was prepared with 0.3 M sodium cacodylate and 6 mM CaCl2, adjusted to pH 7.4. To improve fixation, the tissue was rapidly microwaved in the fixative solution using a microwave system (cat. no. 36700, Ted Pella) equipped with a power controller, steady-temperature water recirculator, and cold spot. The microwaving protocol, based on Tapia et al. (2012)^57^, was performed as follows: at power level 1 (100 W) for 1 min on, 1 min off, 1 min on, followed by power level 3 (300 W) for 20 s on, 20 s off, 20 s on, repeated three times. Fixation was continued overnight at 4°C in the same solution. The following day, the sample was washed three times (30 min each) in 0.5x cacodylate buffer before osmication with 2% OsO4 in 0.5x cacodylate buffer for 90 minutes. After a brief wash in 0.5x cacodylate buffer (<1 min), the sample was incubated in 2.5% potassium ferrocyanide in 0.5x cacodylate buffer for 90 minutes. The sample was then washed with filtered water (three exchanges, 30 min each) before being incubated in 1% (w/v) thiocarbohydrazide (TCH) in filtered water for 45 minutes. Due to poor dissolution of TCH, the solution was preheated at 60°C for ∼90 minutes with occasional shaking, then cooled to RT for 5 minutes prior to incubation. After incubation, the sample was washed with filtered water (three exchanges, 30 min each). A second osmication was performed in 2% OsO4 in filtered water for 90 minutes, followed by three additional washes (30 min each). En-bloc staining was then carried out overnight at 4°C using 1% uranyl acetate in filtered water. The uranyl acetate solution was sonicated for 90 minutes and filtered with a 0.22 μm syringe filter before use. All steps involving uranyl acetate were performed in the dark. The next day, the sample was washed with filtered water (three exchanges, 30 min each) and dehydrated through a graded ethanol series (25%, 50%, 75%, 90%, 100%, and 100% ethanol, 10 minutes each step). The sample was then transferred to 100% propylene oxide (PO) (two exchanges, 30 minutes each). Infiltration was performed with LX112 epoxy resin (Ladd, 21212) mixed with PO in a series of graded steps (25% resin/75% PO, 50% resin/50% PO, 75% resin/25% PO, and 100% resin, with each step lasting 4 hours). The sample was mounted in fresh resin in a mouse brain support tissue [60], with the head exposed to facilitate cutting. The mouse tissue, fixed using standard procedures [62], was cut into 2–3 mm cubes, which were pierced with a 0.75 mm puncher (EMS, 57395) to insert the larva. The cubes were stained alongside the fish samples using the protocol described above, except that the uranyl acetate overnight step was performed at RT. The samples were then cured with support tissue for 3 days at 60°C. During all steps, a rotator was used. Aqueous solutions were prepared with water purified through a filtration system (Arium 611VF, Sartorius Stedim Biotech). The entire protocol, including surgery, fixation, staining, and resin embedding, was completed in 5 consecutive days, followed by 3 days of resin curing. The cured block then was trimmed into a diamond shape. Sections (30–35 nm thick) were automatically collected on carbon-coated tape using a custom tape collection device (ATUM) mounted to a commercial ultramicrotome [60]. The tape was cut into strips and deposited on 30 silicon wafers, which were post-stained as previously described [60]. Wafers were then mounted on the 61-beam multiSEM (MultiSEM 505, ZEISS) stage, and the position of each section was determined using a reflected light microscope to guide high-resolution imaging. Imaging was performed at 4x4 nm pixel resolution using secondary electron emission with a dwell time of 400 ns per pixel. The quality of each individual section was assessed using previously described methods [63]. Cells within a 1 µm-thick slab, positioned posterior to and partially encompassing a cardiac neuron, were manually annotated using VAST8 [61]. Note that this sample was identical to Boulanger-Weill et al.[64], in preparation.

### Custom setup for hemodynamic measurements

Functional imaging for heart rate during looming was conducted using a custom-built optical path based on the Navitar 6000 zoom system. We used a compounded magnification of 8x. To avoid visual artifacts, we used a near-infrared (NIR) ring of LEDs at 750nm and a 750nm band-pass filter (Semrock, 88-013). We acquired 84 frames per second using a Grasshopper 3.0 NIR camera (Teledyne Flir Model: GS3-U3-41C6NIR-C: CMOSIS CMV4000-3E12, NIR). The illumination LEDs were driven using pulse width modulation at 10% duty cycle to avoid heating the sample. The fish was embedded in 2% agarose and then the tail was released using a micro scalpel. This is the same embedding protocol described above under “Drug treatment”. Fish were embedded 1 hour before recordings. Once a fish was embedded, it was placed over a small opening of filter paper that was used to project images from a mini projector. Looming stimuli were designed to be a dot that increased in size and became darker following the derivative of an arc tangent function, which is the equivalent of assuming a predator approaches at constant acceleration. The maximum size of the dot was reached in 6 seconds, and after that, the fish remained in darkness for another 12 seconds before the background light appeared again. To prevent adaptation, fish were exposed to only one stimulus every 5 minutes. Stimuli were repeated 5 times per fish. Similarly, for dark flash experiments background red light was turned off for 32 seconds every five minutes. Tail flicks were tracked using custom Python code previously presented in [65].

### Heart rate quantification

Heart rate was quantified from the periodic near-infrared intensity changes associated with atrial or ventricular contractions. After denoising and detrending the chamber signal, consecutive cardiac cycles were identified and the duration between successive beats was measured. Each instantaneous heart-rate value was calculated as the inverse of the corresponding cardiac interval, yielding a beat-resolved heart-rate trajectory. These beat-derived values were used for the cardiac analyses and as the target or predictor in the neural models described below.

### Analysis of developmental cardiac variability

Heart rate was derived from individual cardiac intervals, with each value in the heart-rate time series calculated as the inverse of the duration of one heartbeat. Variability analyses were performed on the sequence of interbeat intervals rather than directly on the heart-rate values.

For each trial, we analyzed a 60-s pre-stimulus baseline period ending 5 s before stimulus onset. The first stimulus began 268 s after the start of the recording, and subsequent stimulus onsets were separated by 300 s. Baseline periods were analyzed separately so that repeated measurements from the same fish could be retained in the mixed-effects models. Windows containing fewer than 30 valid heartbeats or fewer than 20 valid pairs of consecutive intervals were excluded.

Heart-rate values outside the physiological range of 0.5–8 Hz were excluded. Additional isolated artifacts were identified relative to a 15-beat rolling median and excluded when their deviation exceeded six scaled local median absolute deviations. Successive-difference calculations were performed only within uninterrupted sequences of accepted heartbeats. Thus, RMSSD was not calculated across an excluded or missing interval.

For each baseline period, we calculated the mean interbeat interval, the standard deviation of interbeat intervals, and RMSSD. RMSSD was calculated as

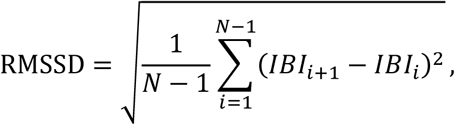

where *IBI_i_*and *IBI_i_*_+1_are consecutive interbeat intervals. RMSSD therefore increases when adjacent cardiac intervals differ more strongly in duration. To compare animals with different mean cardiac intervals, RMSSD was normalized to the mean interbeat interval:

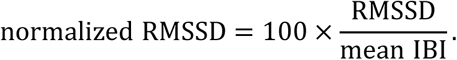

We additionally calculated the coefficient of variation of interbeat intervals, CVNN:

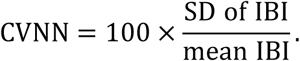

RMSSD and CVNN provide complementary descriptions of cardiac variability. RMSSD emphasizes changes between consecutive cardiac intervals, whereas CVNN measures the overall dispersion of all intervals within the analyzed period. Metrics were calculated separately for each baseline period and then averaged within each fish for visualization and fish-level descriptive statistics.

Developmental differences in fish-level normalized RMSSD and CVNN were evaluated using Kruskal–Wallis tests. These nonparametric analyses treated each fish as one independent biological replicate.

To determine whether normalized RMSSD increased gradually across development or underwent a more discrete developmental transition, we compared four linear mixed-effects models. The models described age as: (1) a continuous linear predictor; (2) a binary transition between 4 and 5 dpf; (3) a transition between 4 and 5 dpf followed by a continuous slope from 5 to 12 dpf; or (4) a categorical developmental-stage variable. All models were fitted to log-transformed normalized RMSSD values from individual baseline periods and included trial number as a fixed effect and fish identity as a random intercept:

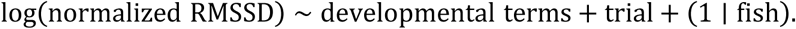

Models were initially fitted by maximum likelihood and compared using the Akaike information criterion, Bayesian information criterion, differences in Akaike information criterion, and Akaike weights. The selected model was refitted using restricted maximum likelihood for coefficient estimation and confidence intervals. Population-level predictions were evaluated at the mean trial number and back-transformed to the original normalized-RMSSD scale. The developmental data were best described by a transition at 5 dpf followed by a more gradual subsequent increase.

The magnitude of the 4-to-5 dpf transition was also estimated independently of the mixed-effects model by bootstrap resampling at the fish level. Fish were resampled with replacement within each age group, and the ratio between the median normalized RMSSD at 5 dpf and the median at 4 dpf was recalculated for 10,000 bootstrap samples. The 2.5th and 97.5th percentiles of the bootstrap distribution were used as the 95% confidence interval.

Because heart-rate variability can depend on average heart rate, we next modeled absolute RMSSD in milliseconds as a function of mean interbeat interval and developmental stage. Absolute RMSSD and mean interbeat interval were log transformed. We compared models containing: (1) mean interbeat interval alone; (2) mean interbeat interval and the piecewise developmental transition; (3) mean interbeat interval and categorical developmental stage; or (4) an interaction between categorical developmental stage and mean interbeat interval. All models included trial number as a fixed effect and fish identity as a random intercept:

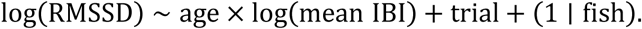

The interaction model received the strongest support by Akaike information criterion, indicating that the relationship between average interbeat interval and cardiac variability differed across developmental stages. Age-specific fitted relationships and population-level 95% confidence intervals were generated from this model with trial number held at its centered value.

### Functional imaging

Similar to the protocol described for embedding fish in the Heart Rate Quantification portion of this Methods section, fish were transferred into the measurement rig after 2 hours of embedding. We used a custom-built two-photon microscope, operated by custom-written Python 3.7-based software [65]. A femtosecond-pulsed MaiTai Ti:Sapphire laser (Spectra Physics) tuned to 920 nm was used to image for GCaMP6s/f. A set of x/y-galvanometers (Cambridge Technology), and a 20× Olympus infrared-optimized objective (XLUMPLFLN) were used to scan over the brain. We collected fluorophore emission using a green photomultiplier tube (PMT), amplified by a current preamplifiers (Stanford SR570). We used frame acquisition rates of around 1.5 Hz in brain imaging experiments. We adjusted laser power to 12.7 mW at the specimen, a low enough value that did not seem to interfere with the physiology of larvae. In our brainstem imaging experiments (Fig. 3), we imaged each plane at a spatial resolution of 0.35 μm per pixel (700 × 700 pixels) for 25 min, while presenting ∼5 trials of 12 s of stimuli interleaved by 5 minutes of rest. We then moved the objective 10 μm to the next imaging plane and repeated the procedure. We acquired 4-6 planes, resulting in a total imaging time of around 1.5 h per fish.

In the case of vagal and sympathetic ganglia, we needed the sagittal view to visualize the entire ganglia, and 2-photon microscopy in this position resulted in slow overheating given the pigmentation in that region.

Because these ganglia are easily accessible with a single-photon microscope from the sagittal view, we used a Zeiss Multiphoton LSM 980 to image these fish. Because the imaging region needed for these ganglia is smaller, we conducted functional imaging at 16.6 Hz. The scanning laser was restricted to the ganglia and therefore did not result in visual artifacts. In these experiments, UV flash at 410nm was used as a visual stressor.

### Optogenetic stimulation

Optogenetic stimulation experiments of the motor limbs of the autonomic system were conducted using galvo mirrors to confine the illumination region of a 488nm laser to small subsets of neurons. The specificity of the activated region depends on (1) the sparseness of the genetic marker, and (2) the targeted region with the laser. Given the precision of the galvo mirrors used to direct the laser the tolerance for spatial illumination is around ∼1.2 µm. The stimulation pulse lasted 1.2 seconds at 75.6% laser power with a 488nm laser and was repeated every 5 minutes. The preparation of larvae before the experiment was analogous to the functional imaging preparation. In the cases of the sympathetic system and cardiomyocytes, heart rate was measured before and after stimulation, but not during stimulation, because of the location of the stimulus pulse. Each fish was tested for 5 trials. Frames were acquired every 60 ms, and during the 1.2s of optogenetic stimulation, the stimulated region was scanned 20 times. Fish were screened for tdTomato (which is co-expressed with CoChR in our transgenic fish) at least 2 hours before the experiments.

For quantification of APG optogenetic responses (Fig. 5O–Q; Supp. Fig. 7), heart rate was averaged over a 12 s baseline immediately before stimulation and a 12 s response window immediately after the stimulation interval. No heart-rate values were included during the stimulation period because concurrent cardiac imaging was not acquired. For each trial, the response was normalized to the pre-stimulation baseline, and trial responses were averaged within fish. For developmental comparisons of APG-evoked responses, trial responses were averaged within each fish and fish-level ΔHR/baseline values were compared across ages using a Kruskal–Wallis test followed by Tukey–Kramer pairwise comparisons of mean ranks. For the 9 dpf propranolol experiment, individual identities linking measurements obtained during propranolol exposure and after washout were not retained. We therefore assessed the robustness of the drug effect using 10,000 plausible random pairings between the 11 propranolol measurements and subsets of 11 of the 17 post-washout measurements. A two-sided Wilcoxon signed-rank test was calculated for each plausible pairing; 91% of random pairings yielded p < 0.05.

## Other Computation and Modeling Methods

This section details preprocessing, generalized linear models (GLMs), kernel estimation, cross-validation, dimensionality reduction for kernel shape, circular phase statistics, clustering, and quantification of optogenetic effects used throughout the study. Unless otherwise stated, analyses were performed in MATLAB and Python (NumPy/SciPy), with custom code available upon publication on our GitHub page.

### Signals, Notation, and Preprocessing

Let *HR(t)* denote the heart-rate time series (Hz) sampled at interval *Δt*; let *ri(t)* be the calcium signal (ΔF/F₀) for neuron *i*. Signals were detrended (second-order polynomial) and z-scored within fish/session.

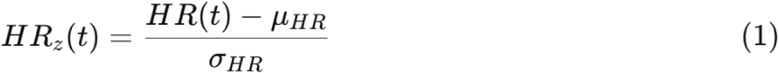

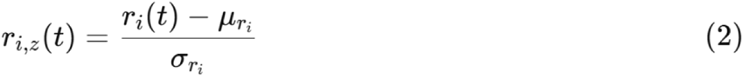

### Causal FIR Design Matrices

For models using discrete time lags, we constructed a lag matrix of HR with causal finite-impulse-response (FIR) supports. For a maximum lag *L* and step δ, the causal HR design matrix *X* was constructed with columns:

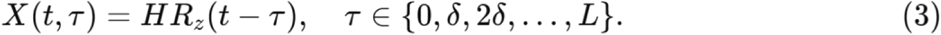

The lag support was model-specific. For the cholinergic hindbrain decoding model (Fig. 3), kernels spanned 0– 4 s in 0.25 s steps. For the APG decoding model (Fig. 5), kernels spanned 0–4 s in 0.1 s steps. Fractional lags were evaluated after temporal interpolation of the input traces.

### GLM for Sensory Encoding (HR → Neural Activity)

Neural activity was modeled as a convolution of HR with a causal kernel *ki(τ)*, plus a bias *bi*.

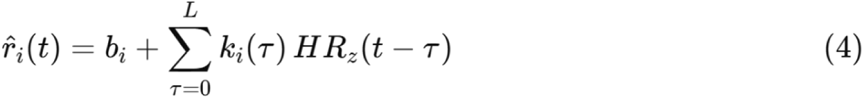

We applied a static nonlinearity for saturation effects:

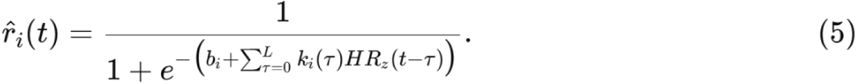

Kernels were estimated by ridge regression:

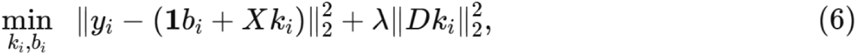

With *D=I* unless otherwise stated, *λ* was chosen by inner cross-validation.

### GLM for Motor Decoding (Neural → HR)

For decoding HR from neural populations:

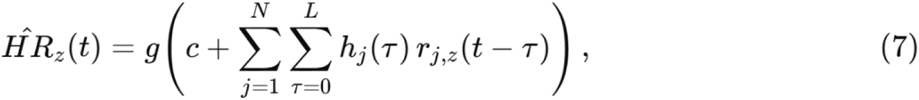

With *g()* a sigmoid as in eq. (5).

### Cross-Validation and Performance

Blocked 5-fold cross-validation: 80% train, 20% test per fold. Model performance was quantified as cross- validated R^2^:

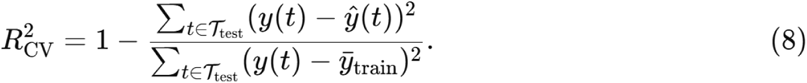

### Kernel Normalization and Shape Metrics

Kernels were L²-normalized for visualization in heatmaps:

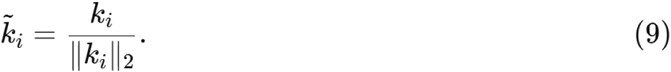

### Kernel PCA and Phase Extraction

Normalized kernels were projected into kernel principal component analysis (PCA) space. Phases were defined from the first two PCs:

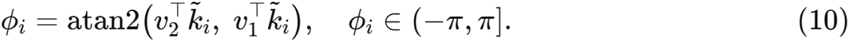

### Circular Statistics and Permutation Tests

For a set of phases {*ϕ_i_*}, the circular mean vector was:

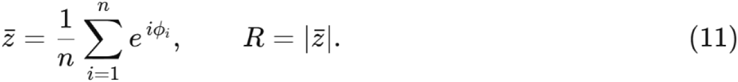

Permutation tests compared observed *R* to a null distribution obtained by shuffling or circularly shifting predictors:

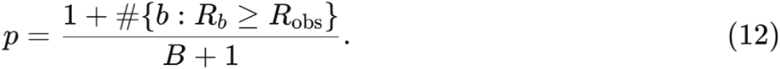

Here the “#{⋅}” notation means “the number of elements satisfying the condition.”

In words:

- R_obs_ is the resultant length of the actual data.
- R_b_ are resultant lengths from each of *B* permutations.
- #{b:R_b_≥R_obs_} counts how many permutation values were greater than or equal to the observed one.

This is the standard way of estimating a permutation p-value (with the +1 in numerator/denominator for finite-sample correction).

For developmental comparisons of kernel phase (Fig. 7K), phase was defined as the angular position of each normalized kernel in the first two principal-component dimensions. Left- and right-side recordings were pooled within developmental stage. For each pair of developmental stages, we calculated the circular distance between their mean phase directions. Statistical significance was assessed by pooling phases from the two developmental stages and randomly reassigning them between groups while preserving the original group sizes. This procedure was repeated 5,000 times. The permutation p value was calculated as (1 + the number of permuted mean-phase differences greater than or equal to the observed difference)/(5,001).

### Determining the number of clusters of kernels

To determine the optimal number of kernel clusters (K) for Fig. 5, we applied three complementary criteria:

1. Silhouette analysis

1. We computed the average silhouette coefficient across neurons for a range of candidate cluster numbers (K = 2–8).
2. The silhouette score measures how similar each neuron’s kernel is to others in its assigned cluster compared to neighboring clusters.
2. Elbow of explained variance criterion.

1. This criterion quantifies the point at which the explained variance vs principal component becomes flat.
3. Gap statistic

1. The gap statistic compares the within-cluster dispersion of the observed data to that of a null distribution generated from randomized (reference) datasets.
2. The optimal K is the smallest number of clusters such that the observed within-cluster dispersion is substantially smaller than expected under the null.
3. This method guards against overfitting to spurious structure.

All three metrics were computed for candidate values of K, and the chosen solution (K = 6) was supported consistently across methods (Supplementary Fig. 6). We further confirmed stability of the clustering by bootstrap resampling (subsampling neurons and repeating PCA+k-means), which yielded reproducible assignments.

---

**Supplementary Figure 1.**
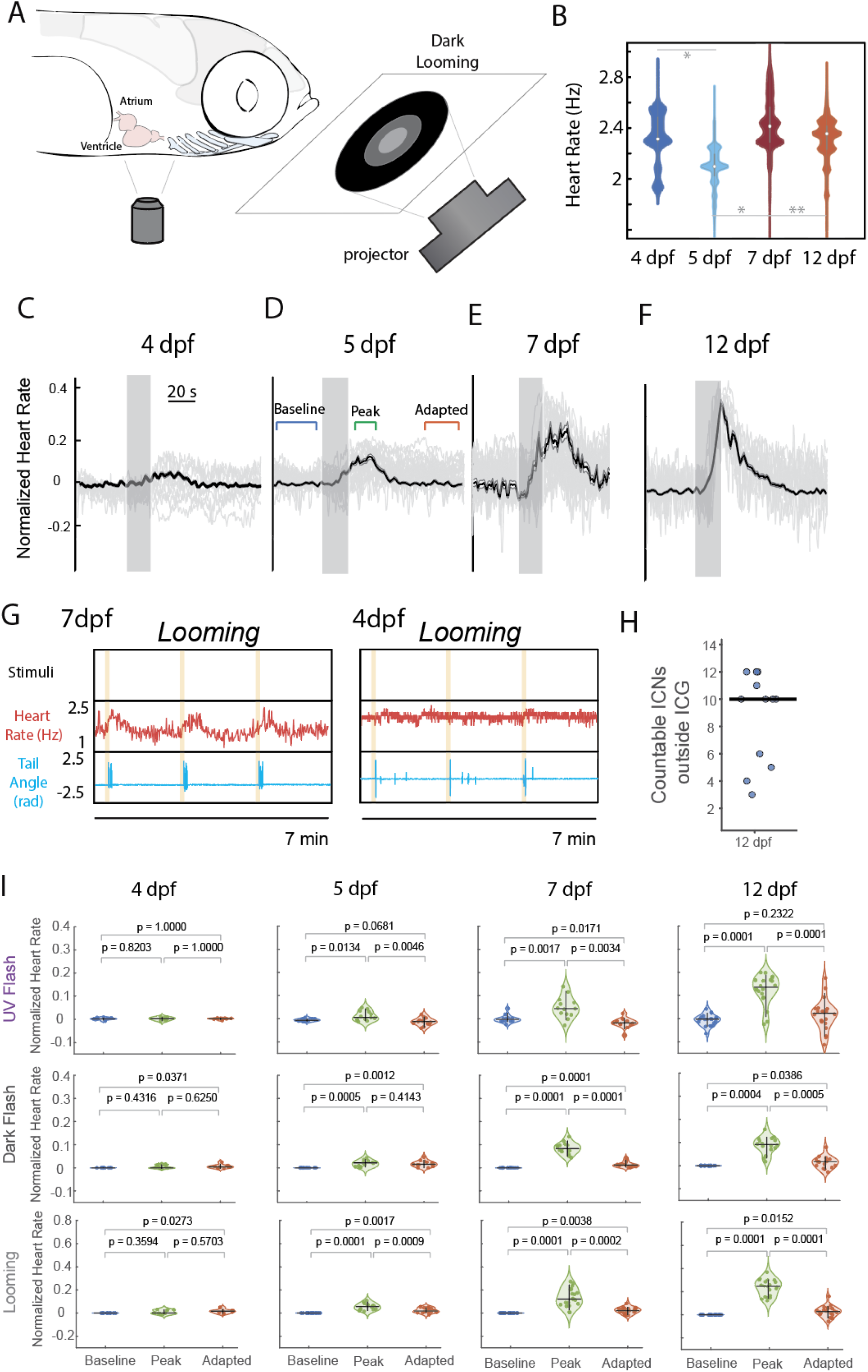
Developmental progression of heart rate responses to visual stimuli. **(A)** Schematic of the experimental setup. Larval zebrafish were embedded with the tail released, and visual looming stimuli were projected from below, while near-infrared imaging was used to extract heart rate. **(B)** Baseline heart rate distributions across development (4, 5, 7, and 12 dpf). Each violin shows the distribution across fish, with median and interquartile ranges indicated (n = 15–20 fish per age). * indicate p<0.0001 and ** indicate p< 0.001. p-values from Steel-Dwass test. **(C–F)** Average heart rate responses to looming stimuli at (C) 4 dpf, (D) 5 dpf, (E) 7 dpf, and (F) 12 dpf. Thin gray traces represent individual animals, the thick black line is the mean across fish, and shaded regions denote the Standard Error (SE) of the mean. Vertical gray bars mark the stimulus presentation. **(G)** Representative simultaneous recordings of heart rate (red) and tail angle (cyan) in response to looming at 7 dpf (left) and 4 dpf (right). At both ages, tail flicks indicate behavioral responses, but only the 7 dpf fish show robust heart rate modulation. **(H)** Quantification of reliably countable ICNs outside the intracardiac ganglion (ICG) at 12 dpf. Each point represents one fish. Counts include ICNs associated with the SAP and AVP, but exclude the ICG because individual somata within the ganglion were frequently overlapping and could not be counted reliably across animals. Horizontal line indicates the median. (I) Quantification of heart rate responses to UV flash, dark flash, and looming across development. Violin plots show normalized heart rate in baseline, peak, and adapted epochs for each fish (5 trials averaged per animal). Each dot represents one fish; horizontal bars denote medians and vertical bars denote the 90% interval range. p-values from the Steel–Dwass test are reported for each pairwise comparison.

**Supplementary Figure 2.**
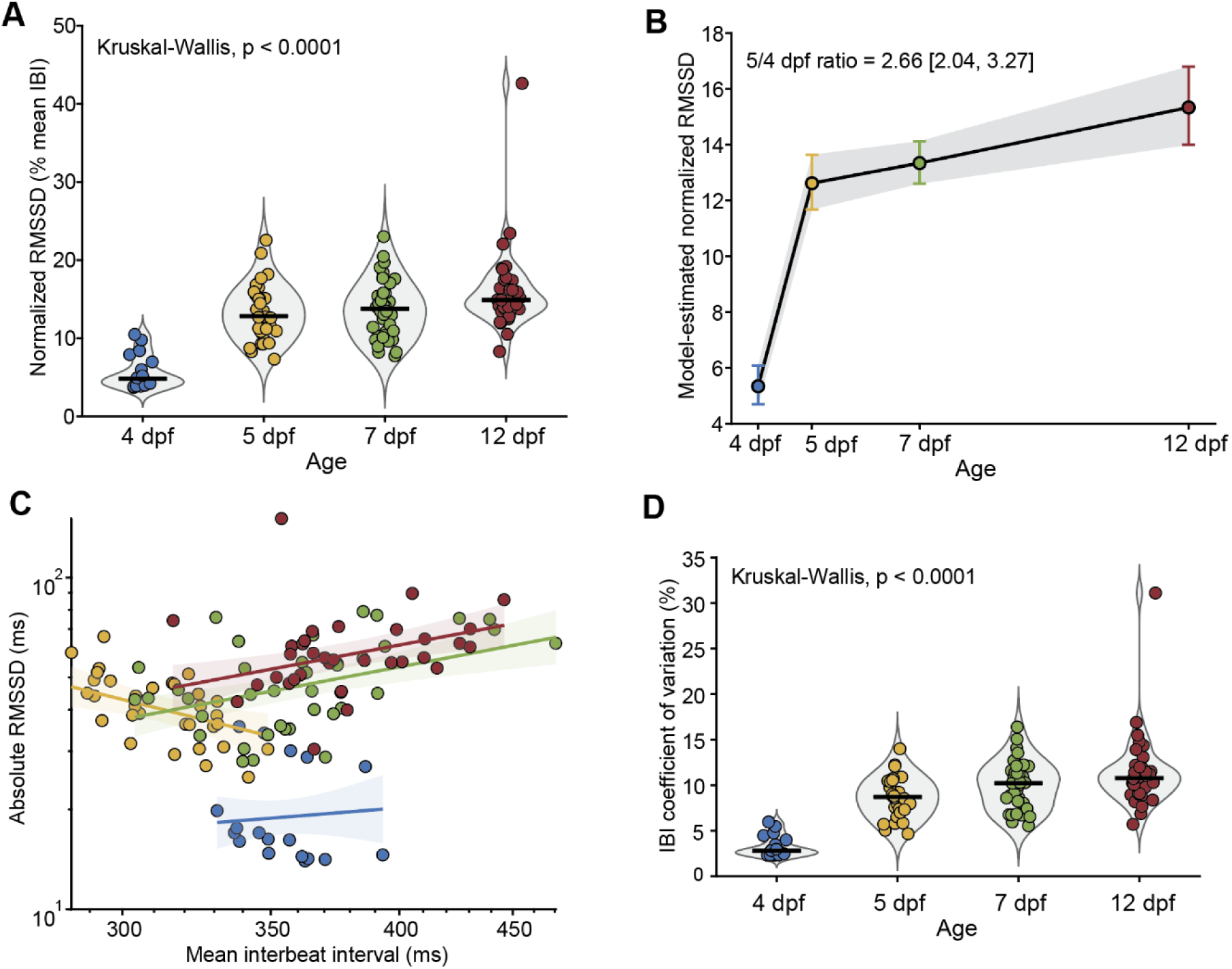
Baseline cardiac variability increases during development. Cardiac variability was quantified from pre-stimulus baseline recordings at 4, 5, 7, and 12 days post-fertilization (dpf). Each cardiac interval was calculated from the duration between consecutive heartbeats. Metrics were first calculated separately for each baseline period and then averaged within each fish, such that each point represents one fish. Colors denote developmental stage throughout: 4 dpf, blue; 5 dpf, yellow; 7 dpf, green; and 12 dpf, red. **(A)** Developmental changes in normalized root mean square of successive differences (RMSSD). RMSSD was divided by the mean interbeat interval and expressed as a percentage. Violin plots show distributions across fish; horizontal black lines indicate medians. Normalized RMSSD differed significantly across ages (Kruskal– Wallis, p<0.0001). **(B)** Population-level estimates from the developmental mixed-effects model. The data were best described by a transition at 5 dpf followed by a more gradual subsequent increase. Circles show model-estimated normalized RMSSD, error bars show 95% confidence intervals, and the gray region connects confidence intervals. Fish-level bootstrap resampling showed that median normalized RMSSD was 2.66-fold higher at 5 dpf than at 4 dpf (95% CI, 2.04–3.27). **(C)** Relationship between absolute RMSSD and mean interbeat interval. Lines show age-specific predictions from the best-supported mixed-effects model, which included an interaction between developmental stage and mean interbeat interval; shaded regions show 95% confidence intervals. **(D)** Developmental changes in CVNN, calculated as the standard deviation of interbeat intervals divided by their mean and expressed as a percentage. CVNN differed significantly across developmental stages (Kruskal–Wallis, p<0.0001).

**Supplementary Figure 3.**
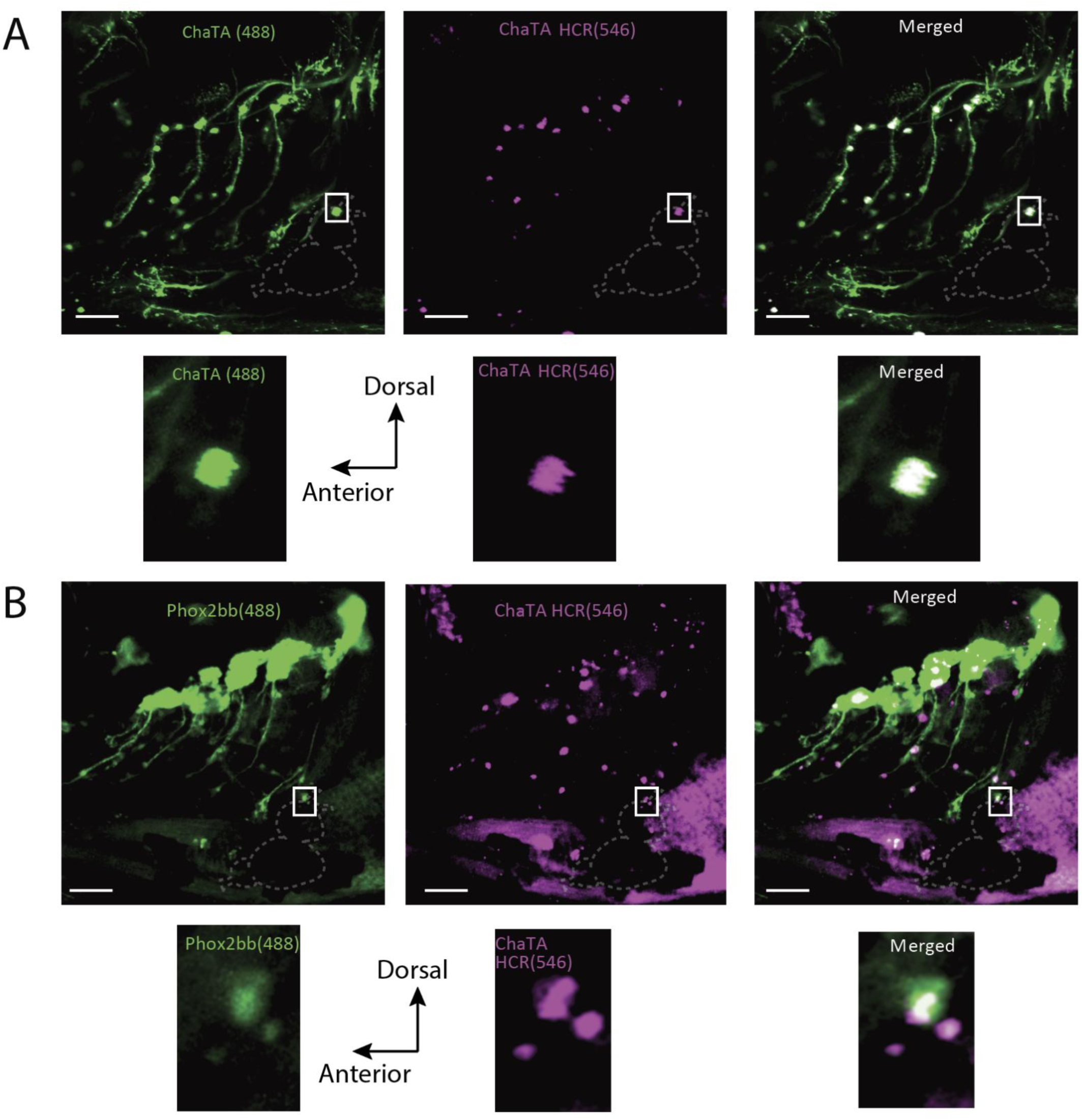
Fluorescent in situ hybridization confirms cholinergic identity of first intracardiac neuron. **(A)** Confocal volumetric imaging of Tg(chata:Gal4, UAS:GFP) larvae. GFP expression (green, 488 nm) labels cholinergic neurons, while hybridization chain reaction (HCR) probes against ChaTA (magenta, 546 nm) confirm transcript expression. Insets show higher-magnification views of a representative neuron (boxed). Merged images demonstrate overlap of GFP and ChaTA HCR signals within intracardiac neurons. **(B)** Confocal volumetric imaging of Tg(phox2bb:EGFP) larvae. GFP expression (green, 488 nm) labels visceral motor and autonomic neurons, while HCR probes against ChaTA (magenta, 546 nm) confirm cholinergic identity. Insets show higher-magnification views of a representative intracardiac neuron (boxed). Merged images again demonstrate colocalization of GFP and ChaTA HCR signals.

**Supplementary Figure 4.**
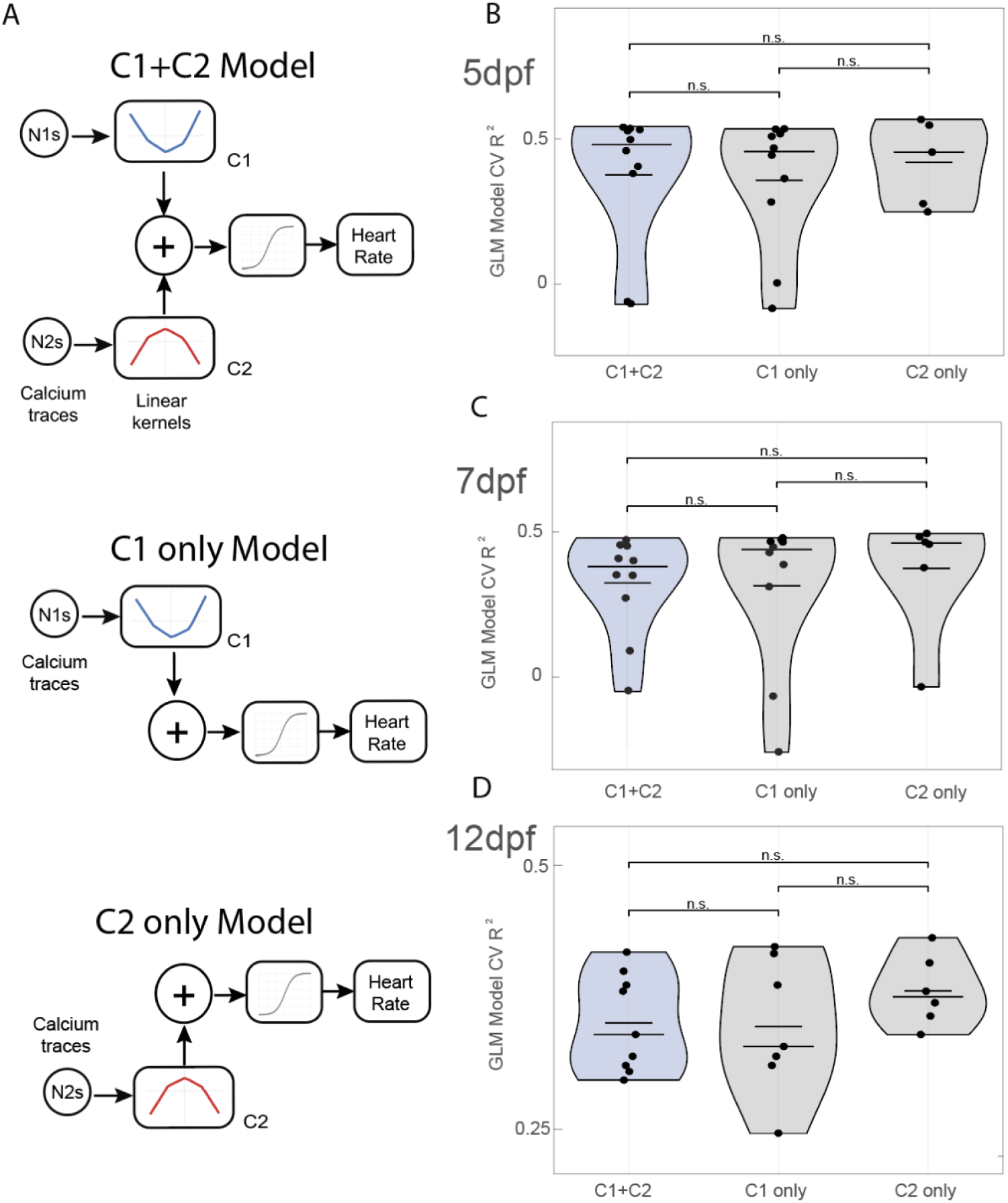
Cluster-restricted GLMs retain heart-rate decoding performance. **(A)** Schematic of the cluster-restricted decoding analysis. The original model used neurons from both kernel classes, C1 and C2, to predict heart rate from cholinergic hindbrain calcium activity. The GLM was retrained using only C1 neurons or only C2 neurons while keeping the same model structure, preprocessing, and cross-validation procedure. **(B–D)** Cross-validated model performance for C1+C2, C1-only, and C2-only models at 5 dpf (B; n=12 fish), 7 dpf (C; n=10 fish), and 12 dpf (D; n=9 fish). Each point represents one fish. Violin plots show distributions across animals; horizontal lines indicate summary statistics. Cluster-restricted models retained substantial predictive performance, with no significant pairwise differences from the combined model at any developmental stage. P value indicates two-sided Wilcoxon signed-rank tests comparing paired fish-level CV R² values between models. n.s. indicates not statistically significant, in all cases p was greater than 0.05.

**Supplementary Figure 5.**
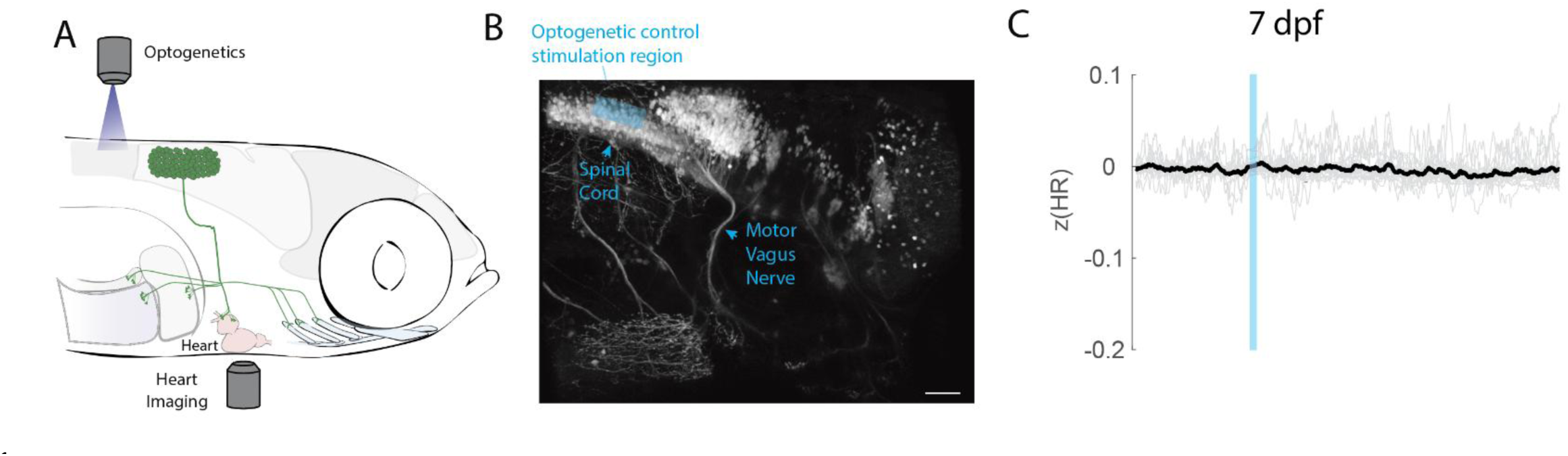
Control optogenetic stimulation of the anterior spinal cord does not affect heart rate. **(A)** Schematic of the experimental setup. Larval zebrafish were embedded for simultaneous heart imaging and targeted optogenetic stimulation. Blue light stimulation was directed to the anterior spinal cord region as a control, outside of the vagal premotor population. **(B)** Confocal image showing the stimulation site (blue shading) relative to the spinal cord and motor vagus nerve. The stimulation region was restricted to the dorsal spinal cord and excluded premotor vagal neurons. Scale bar, 50 μm. **(C)** Heart rate responses at 7 dpf during control spinal cord stimulation. Thin gray traces represent individual trials on 6 fish, the thick black trace shows the mean across fish, and shading denotes the Standard Error of the Mean. No significant change in heart rate was observed, confirming that bradycardic effects in Fig. 3 arise specifically from vagal premotor activation.

**Supplementary Figure 6.**
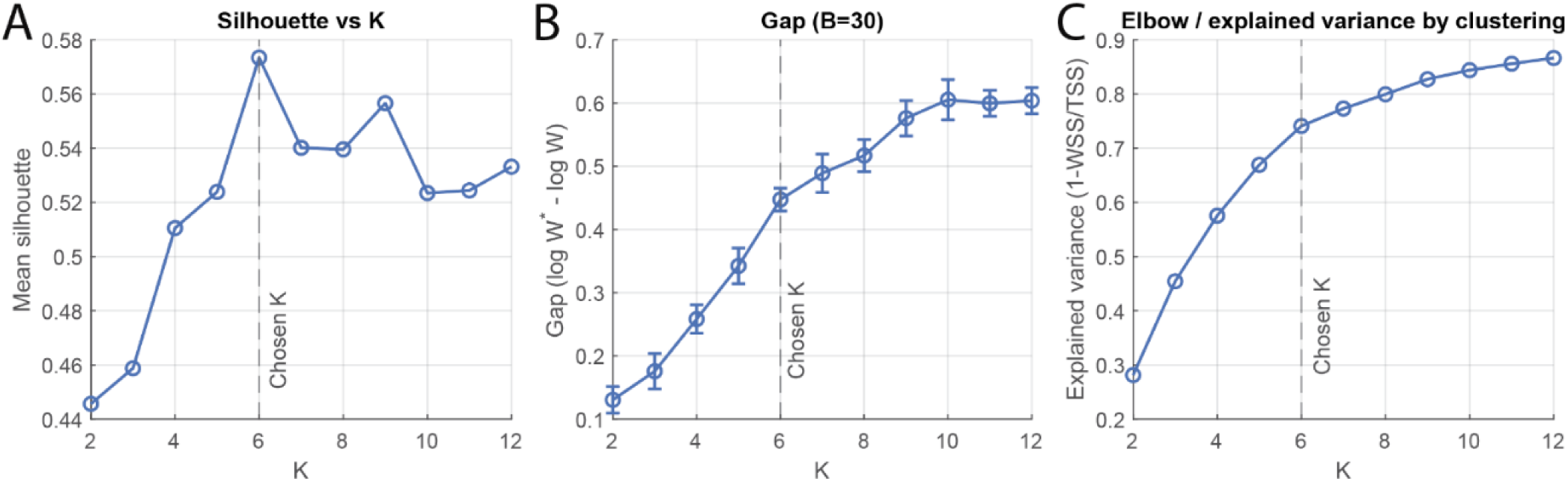
Determining the number of kernel clusters. **(A)** Silhouette analysis for candidate values of K. The silhouette score quantifies the similarity of each kernel to its assigned cluster relative to neighboring clusters. The mean silhouette across neurons is plotted; higher values indicate more coherent clusters. The maximum score was observed at K = 6. **(B)** Gap statistic as a function of K, computed with 30 bootstrap reference datasets (B = 30). The gap statistic measures the separation between observed within-cluster dispersion and that expected under a null distribution. The inflection point at K = 6 supports the chosen cluster number. Error bars denote ± s.e.m. across bootstrap samples. **(C)** Elbow criterion applied to explained variance by clustering. The ratio of between-cluster to total variance increases with K but shows diminishing returns beyond K = 6, indicating this as the optimal choice. Across all three metrics—the silhouette coefficient, gap statistic, and elbow of explained variance—K = 6 was consistently supported as the optimal number of clusters. Bootstrap resampling further confirmed stability of the cluster assignments.

**Supplementary Figure 7.**
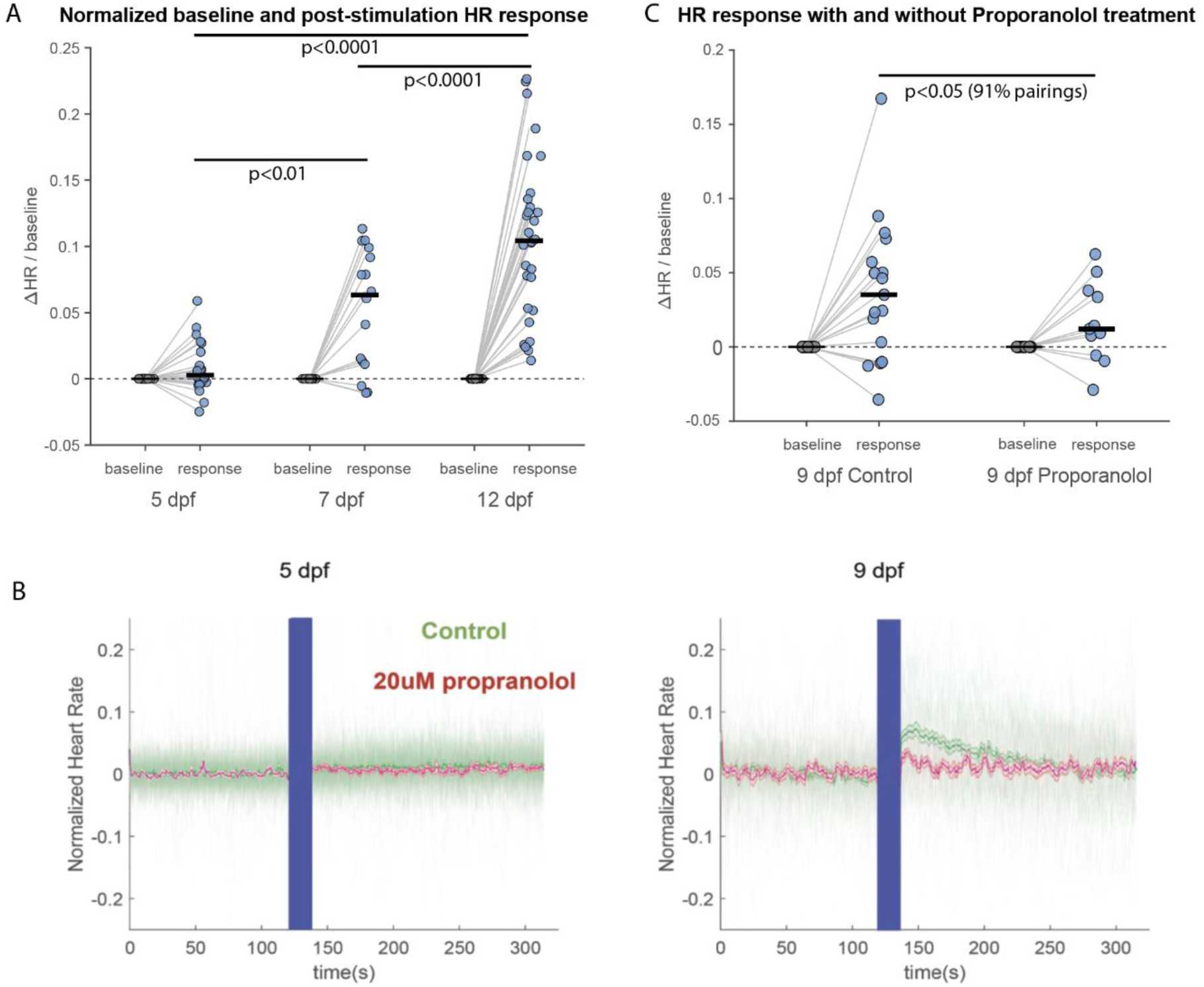
Quantification of APG-evoked cardiac responses. **(A)** Normalized heart-rate responses after APG optogenetic activation at 5 (n=24 fish), 7 (n=16 fish), and 12 dpf (n=30 fish). For each trial, baseline HR was calculated from a 12 s pre-stimulation window and response HR from a 12 s post-stimulation window after excluding the stimulation interval. We normalized responses to baseline and averaged them across trials within each fish. Each point represents one fish; horizontal bars indicate medians. P values indicate pairwise Tukey–Kramer post hoc comparisons of mean ranks following a Kruskal–Wallis test of fish ΔHR/baseline responses across developmental stages. **(B)** Mean normalized heart-rate traces for control and 20 µM propranolol-treated larvae at 5 and 9 dpf. The blue bar indicates optogenetic stimulation, during which concurrent cardiac imaging was not acquired. **(C)** Normalized heart-rate responses during propranolol treatment (n = 11) and after washout (n = 17). Because individual fish identities of the treated population were not tracked, we used a random-matching sensitivity analysis across 10,000 plausible pairings, which showed that 91% of matchings yielded a significant difference at p < 0.05 using a two-sided Wilcoxon signed-rank test.

**Supplementary Figure 8.**
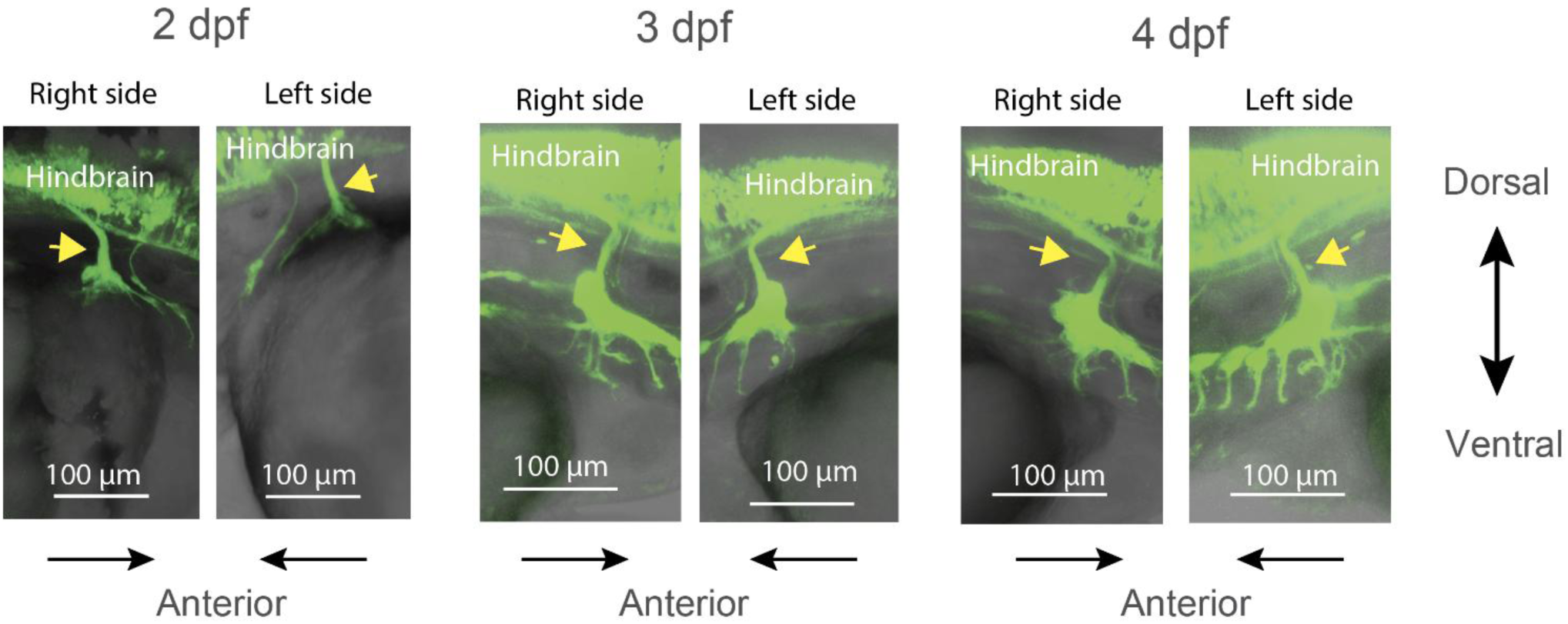
vglut2a-positive vagal sensory projections enter the hindbrain in larval zebrafish. Representative lateral images of the right and left vagal sensory ganglion region at 2, 3, and 4 days post-fertilization (dpf) in vglut2a-Gal4; UAS-GFP larvae. GFP-positive vagal sensory-associated projections extend from the ganglion region into the hindbrain across these developmental stages. Yellow arrowheads mark the central projection. Fluorescence is overlaid on transmitted-light images. Dorsal is up, and anterior is indicated by the arrows below each image. Scale bars, 100 µm. Six animals were examined at each age, and the central projection was observed in all animals.

**Supplementary Figure 9.**
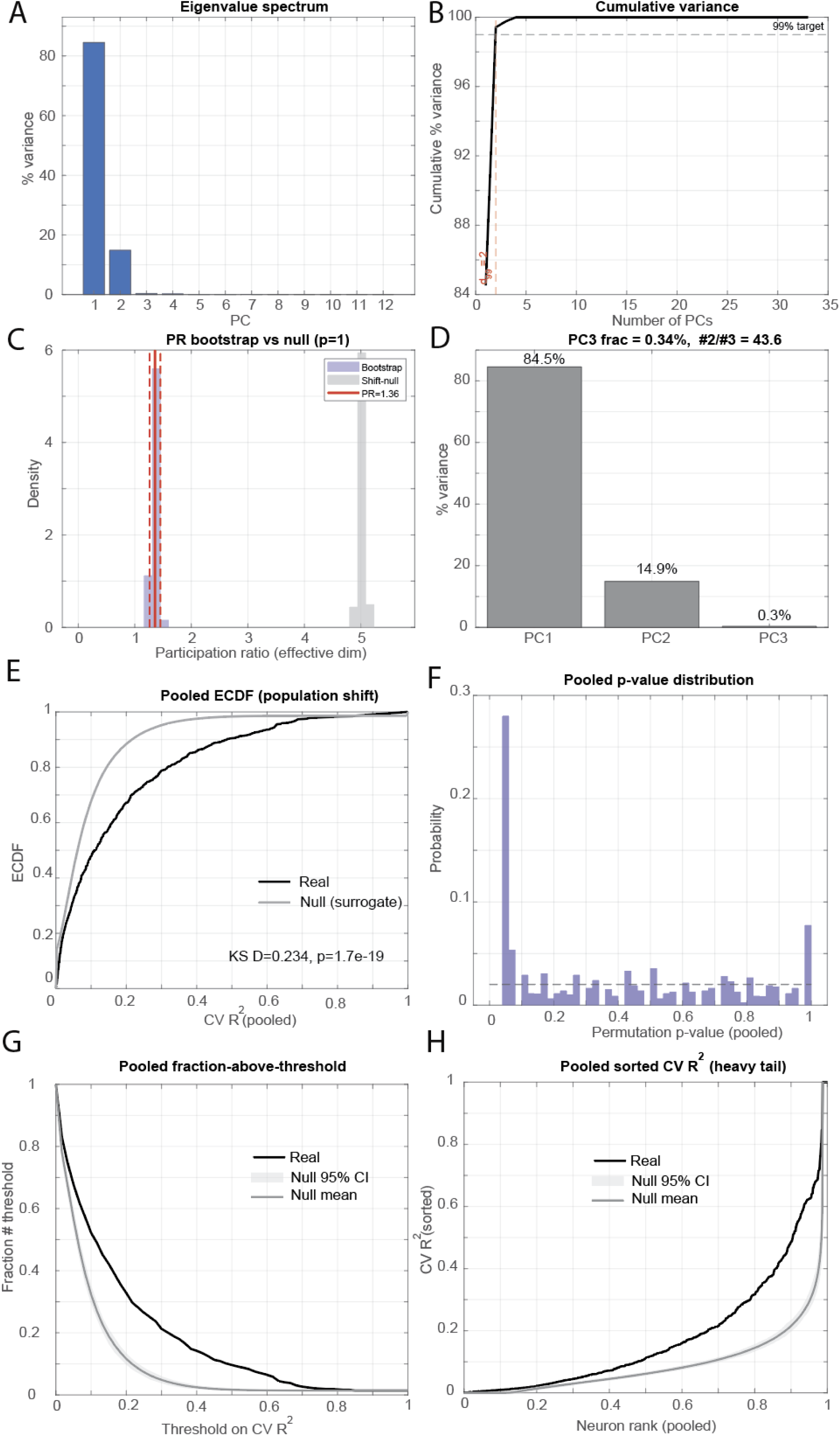
Dimensionality and statistical validation of kernel encoding. **(A)** Eigenvalue spectrum of the kernel covariance matrix. The first two principal components (PCs) capture nearly all of the variance, with PC1 explaining ∼85% and PC2 ∼15%. **(B)** Cumulative variance explained as a function of the number of PCs. A 99% variance target (dashed line) is reached by the first two PCs. **(C)** Participation ratio (PR) analysis comparing bootstrap resampling (purple) with a shift-null distribution (gray). The observed effective dimensionality (red line, PR = 1.36) is far smaller than expected under the null, confirming low-dimensional structure. **(D)** Variance explained by the first three PCs. PC1 accounts for 84.5%, PC2 for 14.9%, and PC3 for only 0.3% of the variance, with PC2 ∼44 times larger than PC3, indicating that kernels are well described by two dimensions. **(E)** Empirical cumulative distribution function (ECDF) of pooled cross-validated prediction performance (CV R2) across neurons. Real data (black) deviate significantly from the surrogate null (gray), Kolmogorov–Smirnov test, D = 0.234, p = 1.7 × 10⁻¹⁹. **(F)** Distribution of permutation p values pooled across neurons. The enrichment of very small p values indicates significant encoding across the population. **(G)** Fraction of neurons above threshold as a function of CV R2. Real data (black) exceed the null mean (light gray) and 95% confidence interval (dark gray shading) across thresholds, demonstrating more neurons with predictive power than expected by chance. **(H)** Sorted CV R2 values across all neurons. The heavy-tailed distribution of real neurons (black) lies above the null distribution (gray), showing that a subset of neurons carries disproportionately strong encoding signals.

## Supplementary Video Legends

Nomenclature:

A:Atrium, V: Ventricle, BA: Bulbous Arteriosus

SAP: SinoAtrial Plexus

AVP: AtrioVentricular Plexus

ICN: IntraCardiac Neuron

ICG: IntraCardiac Ganglion

Supplementary video 1: Example of a 5 dpf *phox2bb-GFP* zebrafish. We identify neurites in the SAP.

Supplementary video 2: Example of a 7 dpf *phox2bb-GFP* zebrafish. We identify the first ICN.

Supplementary video 3: Example of a 12 dpf *phox2bb-GFP* zebrafish. At this age, the ICG has appeared in the atrium, and more neurons are in the SAP and AVP.

Supplementary video 4: Example of a 4 dpf *ChaTA-Gal4, UAS-GFP* larval zebrafish. At this age the cardiac branch of the motor vagus nerve is not yet developed, thus there are no neurites in the recording of the beating heart.

Supplementary video 5: Example of a 5 dpf *ChaTA-Gal4, UAS-GFP* larval zebrafish. At this age the cardiac branch of the motor vagus nerve has reached the heart and it can be seen moving with the SAP during atrium contractions (yellow arrow in the video).

Supplementary video 6: Example of a 7 dpf *ChaTA-Gal4, UAS-GFP* zebrafish. At this age the first ICNS neuron has reached the heart, and it moves with atrium contractions.

Supplementary video 7: Example of a 12 dpf *ChaTA-Gal4, UAS-GFP* zebrafish. At this age the neurites have expanded to reach the AVP (arrow in the video).

